# Resolving marine–freshwater transitions by diatoms through a fog of discordant gene trees

**DOI:** 10.1101/2022.08.12.503770

**Authors:** Wade R. Roberts, Elizabeth C. Ruck, Kala M. Downey, Eveline Pinseel, Andrew J. Alverson

**Author notes:** Corresponding author: Andrew J. Alverson, Department of Biological Sciences, University of Arkansas, 1 University of Arkansas, Fayetteville, AR, 72701, USA.

## Abstract

Despite the obstacles facing marine colonists, most lineages of aquatic organisms have colonized and diversified in freshwaters repeatedly. These transitions can trigger rapid morphological or physiological change and, on longer timescales, lead to increased rates of speciation and extinction. Diatoms are a lineage of ancestrally marine microalgae that have diversified throughout freshwater habitats worldwide. We generated a phylogenomic dataset of genomes and transcriptomes for 59 diatom taxa to resolve freshwater transitions in one lineage, the Thalassiosirales. Although most parts of the species tree were consistently resolved with strong support, we had difficulties resolving a Paleocene radiation, which affected the placement of one freshwater lineage. This and other parts of the tree were characterized by high levels of gene tree discordance caused by incomplete lineage sorting and low phylogenetic signal. Despite differences in species trees inferred from concatenation versus summary methods and codons versus amino acids, traditional methods of ancestral state reconstruction supported six transitions into freshwaters, two of which led to subsequent species diversification. Evidence from gene trees, protein alignments, and diatom life history together suggest that habitat transitions were largely the product of homoplasy rather than hemiplasy, a condition where transitions occur on branches in gene trees not shared with the species tree. Nevertheless, we identified a small set of putatively hemiplasious genes, many of which have been associated with shifts to low salinity, indicating that hemiplasy played a small but potentially important role in freshwater adaptation. Accounting for differences in evolutionary outcomes, in which some taxa became locked into freshwaters while others were able to return to the ocean or become salinity generalists, might help further distinguish different sources of adaptive mutation in freshwater diatoms.

## INTRODUCTION

From bacteria to animals, the salinity gradient separating marine and freshwater environments poses a significant barrier to the distributions of many organisms (Lozupone and Knight 2007; McCairns and Bernatchez 2010; Kenny et al. 2019). Identifying how different lineages cross the salinity divide will improve our understanding of lineage diversification (Dittami et al. 2017) and the adaptive potential of species to climate change (Dickson et al. 2002; Lee et al. 2022). Diatoms are a diverse lineage of microalgae that occur throughout marine and freshwaters, and despite the numerous obstacles facing marine colonists (Kirst 1990, 1996; Nakov et al. 2020), ancestrally marine diatoms have successfully colonized and diversified in freshwaters repeatedly throughout their history (Nakov et al. 2019). These patterns are based on phylogenetic analyses of a small number of molecular markers, however, so they lack the insights of phylogenomic approaches, which can resolve large-scale macroevolutionary patterns and, at the same time, uncover key processes at play during important evolutionary transitions (Pease et al. 2016).

Although phylogenomic datasets have helped resolve historically recalcitrant nodes across the tree of life, they have also revealed how discordance in the evolutionary histories of different genes can confound inferences of species relationships. Systematic errors that lead to gene tree discordance can be caused by biological sources, such as incomplete lineage sorting (ILS) and hybridization (Maddison 1997; Degnan and Rosenberg 2006a), or methodological sources, such as character sampling, compositional heterogeneity, and gene tree error (Foster 2004; Philippe et al. 2011; Xi et al. 2015; Molloy and Warnow 2018). Each of these challenges our ability to resolve species relationships and impacts downstream analyses, such as estimation of divergence times (Smith et al. 2018). Multiple strategies have been proposed to overcome different sources of error, such as excluding third-codon positions from DNA datasets (Sanderson et al. 2000), using site-heterogeneous models for amino acid data (Wang et al. 2018), and identifying the conditions under which concatenation (Edwards 2009) or gene tree summary approaches (Mirarab et al. 2014; Liu et al. 2015) more accurately resolve species relationships.

Discordance between gene and species trees can confound inferences of trait evolution as well (Hahn and Nakhleh 2016), particularly for complex traits that appear to have evolved convergently. Focus on the species tree alone, without considering discordant gene trees, can lead to artifactual inferences of molecular convergence (Mendes et al. 2016). This failure occurs when a trait is determined by genes with tree topologies that do not match the species topology, a condition known as hemiplasy (Avise and Robinson 2008; Hahn and Nakhleh 2016; Storz 2016). Hemiplasy has been identified as a likely explanation for patterns of character incongruence in amino acid substitutions in columnar cacti (Copetti et al. 2017), flower and fruit traits in *Jaltomata* (Solanaceae) (Wu et al. 2018), and dietary specialization in *Dysdera* spiders (Vizueta et al. 2019). In conditions where decreased gene concordance is coupled with short branch lengths and shallow time scales, hemiplasy is expected to have a higher impact on trait reconstruction (Hahn and Nakhleh 2016).

Adaptation to low salinity is a complex trait, so the genomic changes associated with successful freshwater colonizations are multifaceted (Artemov et al. 2017; Cabello-Yeves and Rodriguez-Valera 2019; Rogers et al. 2021) and generally involve mutations in multiple genes and pathways (Jones et al. 2012; Terekhanova et al. 2019; Chen et al. 2021). To better understand the pattern, timing, and process of marine–freshwater transitions by diatoms, we assembled a dataset of 45 genomes and 42 transcriptomes—most of them newly sequenced—to resolve species relationships, explore the causes and consequences of gene tree discordance, and provide insight into how discordance impacted trait reconstruction.

## MATERIALS & METHODS

Detailed methods are provided in Supplementary File S1. Briefly, diatom cultures were isolated from natural plankton or acquired from the National Center for Marine Algae and Microbiota (NCMA) or Roscoff Culture Collection (RCC). Collection data, culture conditions, and voucher information are available in Supplementary Table S1. For genome and transcriptome sequencing, we extracted total DNA and RNA from diatom cultures, constructed sequencing libraries, and sequenced them on the Illumina platform. Based on a large multigene phylogeny of diatoms (Nakov et al. 2018), we included *Coscinodiscus,* Lithodesmiales, and *Eunotogramma* as outgroups. Accession numbers for reads and assemblies are provided in Supplementary Table S2.

We used OrthoFinder (Emms and Kelly 2019) to cluster amino acid sequences from all genomes and transcriptomes into orthogroups, then aligned orthogroups containing ≥20% of the taxa with MAFFT (Katoh and Standley 2013). For each alignment, we identified the best-fit substitution model using ModelFinder (Kalyaanamoorthy et al. 2017) and estimated gene trees with IQ-TREE (Minh et al. 2020b) or FastTree (Price et al. 2010). We then filtered and trimmed the gene trees using the Rooted Ingroup method to produce final ortholog sets (Yang and Smith 2014). This filtering and trimming procedure was performed twice. We used PAL2NAL (Suyama et al. 2006) to reconcile nucleotide coding sequence (CDS) alignments against amino acid alignments. To account for possible saturation at third-codon positions, we removed them from the CDS alignments and, separately, used Degen (Regier et al. 2010; Zwick et al. 2012) to recode synonymous sites in the full coding sequence alignments with the nucleotide ambiguity codes. For example, all leucine codons (CTN, TTR) were degenerated and replaced with YTN. In total, we analyzed datasets consisting of amino acids (AA), first and second codon positions (CDS12), and degenerate codons (DEGEN). We generated final ortholog alignments and inferred trees as described above, using 1000 ultrafast bootstrap replicates to estimate branch support (Minh et al. 2013). We generated summary statistics for final alignments and gene trees with AMAS (Borowiec 2016) and PhyKit (Steenwyk et al. 2021) (Supplementary Table S3). Correspondence analysis of amino acid frequencies across all taxa was performed using GCUA (McInerney 1998).

Species trees were inferred using gene-partitioned maximum likelihood analysis of a concatenated supermatrix with IQ-TREE and the summary quartet approach implemented in ASTRAL-III (Zhang et al. 2018). We performed the matched-pairs test of symmetry in IQ-TREE to identify and remove gene partitions that violated assumptions of stationarity, reversibility, and homogeneity (SRH; Naser-Khdour et al. 2019). Briefly, stationarity refers to the assumption of constant amino acid or nucleotide frequencies across time, reversibility implies that substitutions between amino acids or nucleotides occur equally, and homogeneity implies that the instantaneous substitution rates are constant along a tree or branch (Felsenstein 2004). For the IQ-TREE analysis, we partitioned supermatrices by gene, used ModelFinder to select the best-fit substitution model for each partition, and estimated branch support with 10,000 ultrafast bootstrap replicates. For ASTRAL, we used the ortholog trees as input, collapsing branches with low bootstrap support (<33) to help mitigate gene tree error (Sayyari and Mirarab 2016; Simmons and Gatesy 2021). We estimated branch support using local posterior probability (Sayyari and Mirarab 2016). In addition to heterogeneity in gene histories, compositional heterogeneity can also make species tree inferences difficult at deep time scales (Lartillot and Philippe 2004). To account for this possibility, we estimated a species tree from the concatenated amino acid matrix using the Posterior Mean Site Frequency (PMSF) model implemented in IQ-TREE (Wang et al. 2018).

We calculated the Robinson-Foulds distance between each pair of species trees and visualized the results with a multidimensional scaling plot made with the R package *treespace* (Jombart et al. 2017). We characterized discordance using gene and site concordance factors (Minh et al. 2020a) and quartet concordance factors (Pease et al. 2018). We tested the support for competing backbone topologies in our species tree using the approximately unbiased (AU) test (Shimodaira 2002) implemented in IQ-TREE. Relative gene tree support for the same set of backbone topologies was further evaluated using gene genealogy interrogation method (Arcila et al. 2017). Finally, we performed the polytomy test on the ASTRAL species tree to test whether any of the unstable backbone branches were better represented as a polytomy (Sayyari and Mirarab 2018).

Divergence times were estimated using MCMCtree (Yang 2007; Reis and Yang 2011), using five fossil calibrations (Supplemental File S1), autocorrelated rates, and the approximate likelihood approach. We performed marginal ancestral state reconstruction for marine and freshwater habitat across the species tree using hidden state speciation and extinction models in the R package *hisse* (Beaulieu and O’Meara 2016). Lastly, we explored the potential for hemiplasy in our habitat reconstructions by calculating: (1) hemiplasy risk factors using the R package *pepo* (Guerrero and Hahn 2018); (2) the differences between the number of marine– freshwater transitions optimized on unconstrained versus species-tree-constrained gene trees; and (3) the number of alignment site changes that differed between unconstrained and species-tree-constrained gene trees.

Phylogenetic trees were plotted using the R Bioconductor packages *ggtree* and *treeio* (Yu et al. 2017; Wang et al. 2019). Assembled genomes and transcriptomes have been deposited at NCBI under BioProject PRJNA825288. Supplementary Files, Figures, and Tables are available from the Dryad Digital Repository: https://doi.org/10.5061/dryad.7m0cfxpxp. Voucher images, proteomes, alignments, trees, log files, and code have been deposited in Zenodo: https://doi.org/10.5281/zenodo.7713227.

## RESULTS

We combined 42 newly sequenced draft genomes and 50 newly sequenced transcriptomes with publicly available genome or transcriptome data for a final dataset of 87 taxa representing 59 species. A total of 17 transcriptomes were used directly in phylogenomic analyses, and the other 33 were used for genome annotation. From the combined dataset of genomes and transcriptomes, we generated alignments and gene trees for 6262 orthologs, with each taxon represented in an average of 3275 (52%) orthologs.

### Compositional heterogeneity and dataset construction

Relative Composition Variability (Phillips and Penny 2003) indicated greater compositional heterogeneity in third codon positions compared to amino acids and first and second codon positions (Supplementary Fig. S1). We also found substantial variation in GC content of third codon positions across taxa and genes (Supplementary Fig. S2), ranging from an average proportion of 0.40 ± 0.07 in *Cyclotella kingstonii* to 0.76 ± 0.14 in *Shionodiscus oestrupii*. In addition, nucleotide alignments including all three codon positions had significantly higher levels of saturation compared to the amino acids (Supplementary Fig. S3). To minimize potentially misleading signal in the nucleotide data, we removed third codon positions and, to examine the effects of saturation and GC heterogeneity in coding sequences (CDS), we replaced synonymous signal by recoding CDS sequences with degenerate codons (Zwick et al. 2012). Based on these results, we created three datasets to estimate phylogenetic relationships: amino acids (AA), first and second codon positions (CDS12), and degenerate codons (DEGEN).

Dataset and alignment characteristics are summarized in Table 1 and detailed in Supplementary Table S3. Each dataset initially contained 6262 orthologs (Table 1). Gene trees constructed from all datasets were well supported (80 ± 7% average bootstrap; Table 1; Supplementary Table S3). To reduce systematic errors due to model misspecification, we removed orthologs that failed SRH assumptions (*P* < 0.05), which retained 5522 (AA), 3259 (CDS12), and 3788 (DEGEN) genes in the datasets (hereafter referred to as the “complete” datasets; Table 1). We then subsetted the complete datasets to maximize phylogenetic signal or minimize missing data. To maximize signal and reduce stochastic error, we sorted orthologs by the percentage of parsimony informative (PI) sites and retained the top 25% ranked orthologs (“top-PI”; Table 1). To minimize the amount of missing data and maximize taxon occupancy, we sorted complete datasets by the number of taxa and subsetted these to include the top 25% ranked orthologs with highest taxon occupancy (“top-Taxa”; Table 1). Orthologs in the top-Taxa datasets contained an average of 76 ± 8% of the total taxa.

**Table 1.**
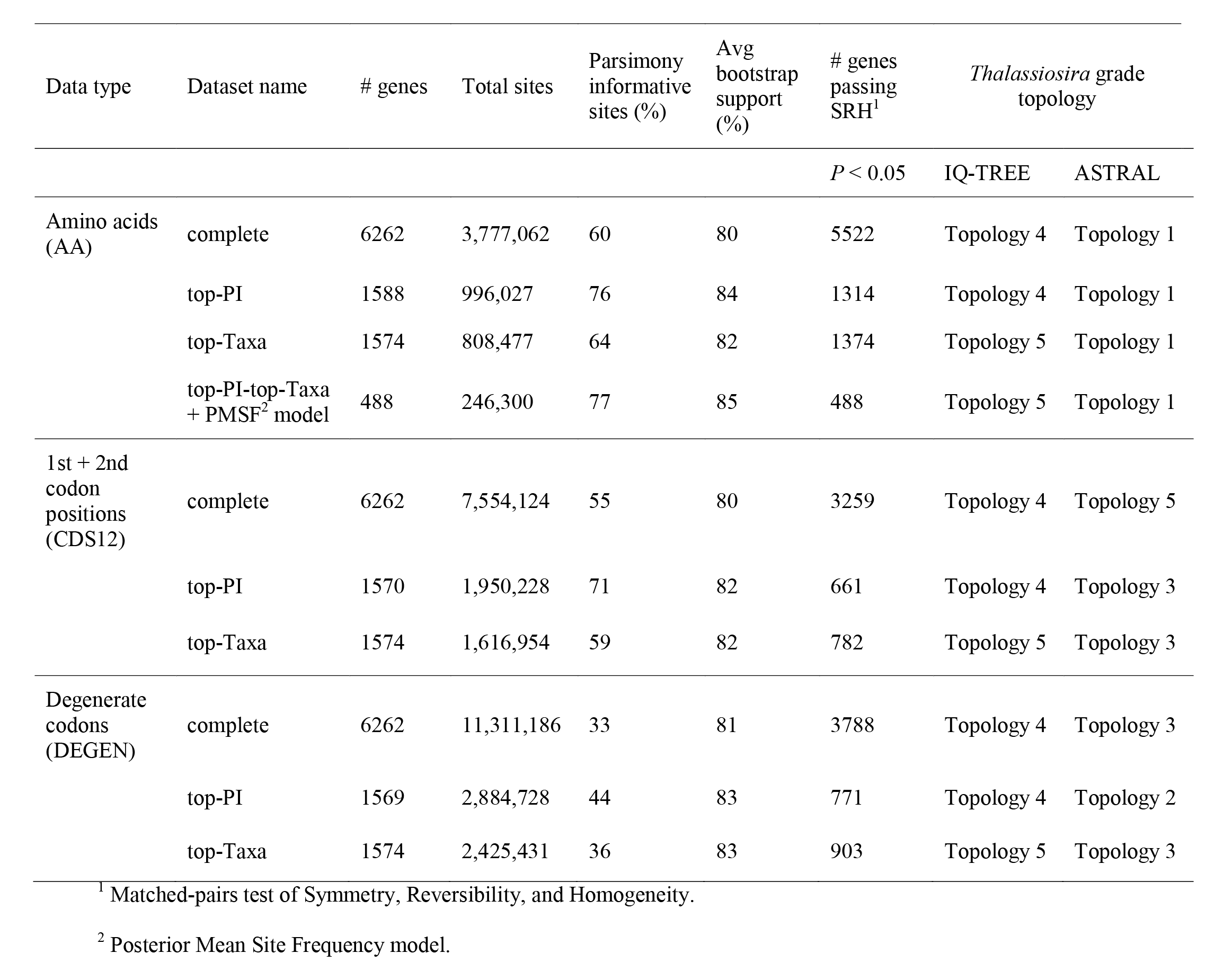
Summary of datasets used in the study.

### Species tree inference and placement of freshwater clades

We initially estimated 18 species trees using amino acids, codon positions 1 and 2, or degenerate codon sequences, with different cutoffs for taxon occupancy or proportion of parsimony informative sites and using both concatenation and summary quartet approaches (Table 1). Correspondence analysis of amino acid frequencies separated taxa principally by habitat (marine versus freshwater) rather than phylogeny (Supplementary Fig. S4), which led us to explore whether compositional heterogeneity affected phylogenetic reconstructions. To do so, we estimated an additional species tree using the PMSF mixture model, which can accommodate heterogeneity in amino acid composition between species at a particular site (Wang et al. 2018). Due to the computational demands of implementing this model, we applied it only to a reduced amino acid dataset with orthologs that met both the top-PI and top-Taxa filtering criteria (“AA- top-PI-top-Taxa”; Table 1).

Previous phylogenetic analyses of this group resolved freshwater taxa into two main clades: the genus *Cyclotella*, which also includes several marine and brackish species, and the “cyclostephanoids”, comprised of several stenohaline genera confined exclusively to freshwaters (Alverson et al. 2011). Given the potential implications for uncovering the mechanisms of freshwater adaptation, we were primarily interested in the placements of these two clades. Gross differences among data types and methods were evident in an ordination of species trees based on pairwise Robinson-Foulds distances, which showed a clear separation between IQ-TREE and ASTRAL topologies, with further separation of IQ-TREE trees estimated from datasets that maximized signal (top-PI) or minimized missing data (top-Taxa) (Supplementary Fig. S5). The phylogenetic position of *Cyclotella* was strongly supported and robust to differences in data type (codons versus amino acids) and analysis (IQ-TREE versus ASTRAL) (Fig. 1). The cyclostephanoids were placed consistently within a large clade of marine *Thalassiosira* and relatives (Fig. 1), but the arrangements of five main subclades—*Thalassiosira* I–IV and the freshwater cyclostephanoids—varied depending on data type and analysis (Table 1; Fig. 2a). We refer to this part of the tree as the *Thalassiosira* grade.

**Figure 1.**
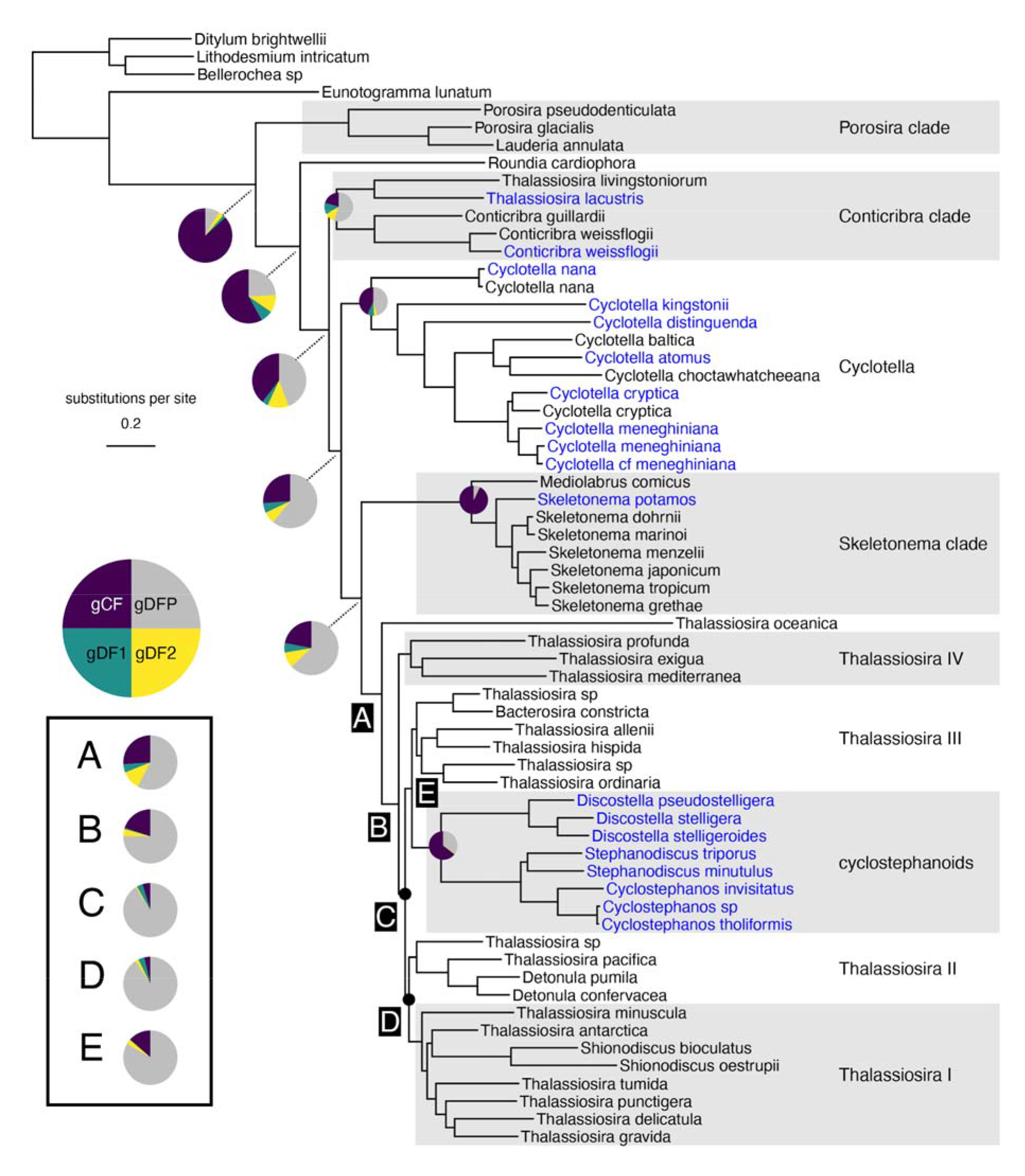
Phylogram based on maximum likelihood analysis of amino acids using the posterior mean site frequency (PMSF) model and a dataset of 488 loci with the highest proportions of taxa and informative sites (“AA-top-PI-top-Taxa” dataset; Table 1). Backbone nodes of the *Thalassiosira* grade are indicated by the letters A–E. All branches had bootstrap support (BS) values of 100 except for those with black circles which had BS = 90. Pie charts on backbone nodes show the proportion of gene trees that support the clade (gCF), the proportion that support both discordant topologies (gDF1, gDF2), and the proportion that are discordant due to paraphyly (gDFP). Size of the pie charts is for clarity only.

**Figure 2.**
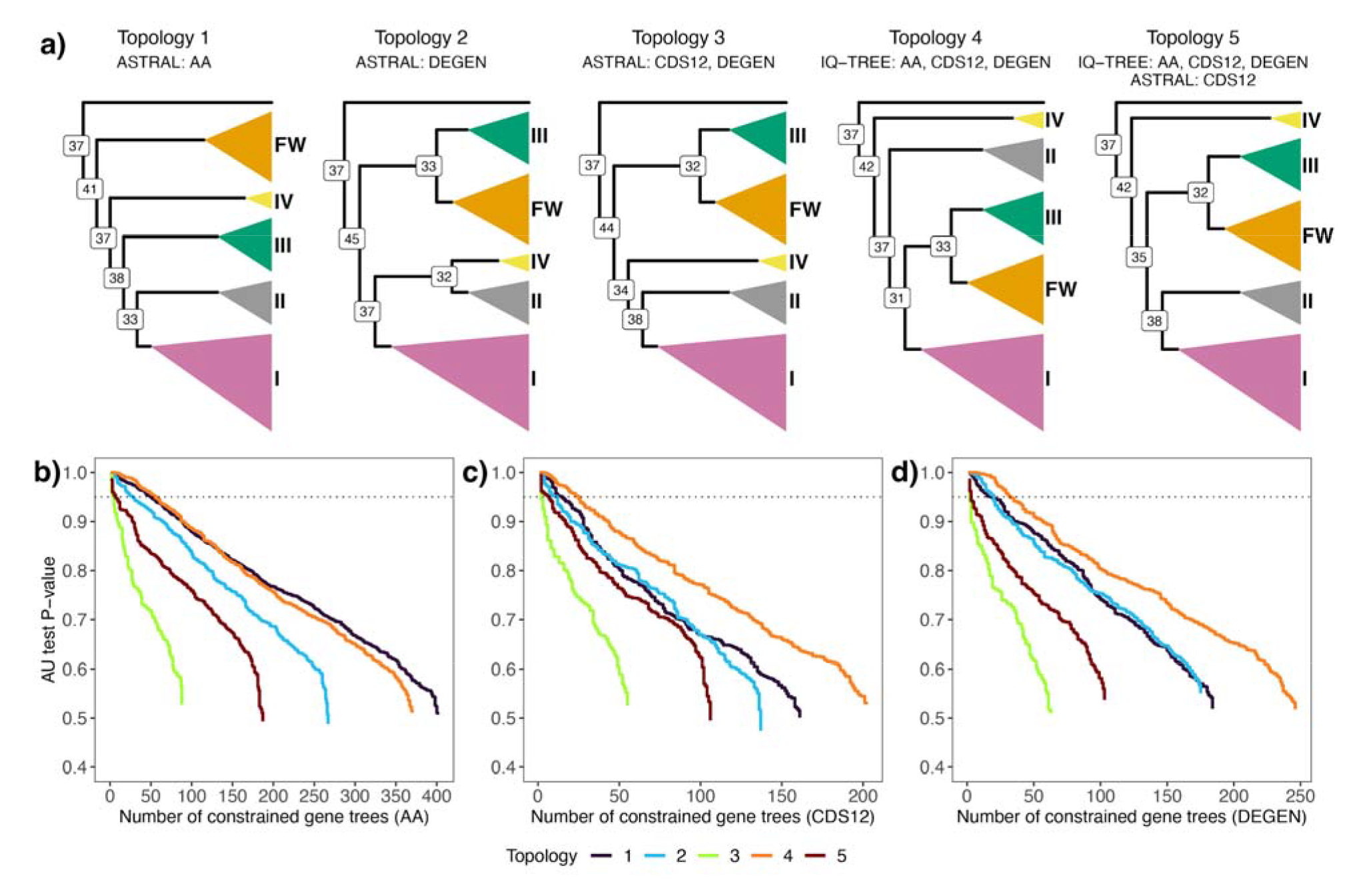
**a)** Phylogenetic hypotheses of the *Thalassiosira* grade inferred using concatenation and summary methods on the amino acid (AA), codon positions 1 and 2 (CDS12), and recoded codon (DEGEN) datasets. Nodes are labeled with the percentage of amino acid sites concordant with the branch (site concordance factor). The principal clade of interest, the freshwater cyclostephanoids, is colored orange and labeled ‘FW’. The four focal clades of marine *Thalassiosira* and allies are labeled I–IV. Panels b–d show results of gene genealogy interrogation tests of alternative hypotheses of relationships within the *Thalassiosira* grade. These tests used datasets filtered to include only the top 25% of orthologs based on the percentage of parsimony informative sites (top-PI) for **b)** amino acids, **c)** codon positions 1 and 2, and **d)** recoded codons. Lines correspond to the cumulative number of genes (x axis) supporting topology hypotheses with the highest probability and their *P* values (y axis) from the Approximately Unbiased (AU) topology tests. Values above the dashed line indicate topological hypotheses that are significantly better than the alternatives (*P* < 0.05). For example, the green line in panel b shows that there were a total of 88 genes that best supported topology 3, though only four of those genes were above the dotted line and were significantly better supported than the other four alternative topologies.

One resolution of the *Thalassiosira* grade (topology 1) was recovered only by ASTRAL analysis of amino acid gene trees and placed the freshwater cyclostephanoids as sister to a clade of *Thalassiosira* I–IV (Table 1; Fig. 2a). All other species trees placed cyclostephanoids as sister to *Thalassiosira* III (Table 1; Fig. 2a). ASTRAL analyses of codon-based gene trees (CDS12 and DEGEN) alone recovered topology 3, which placed cyclostephanoids and *Thalassiosira* III as sister to the remaining *Thalassiosira* (Table 1; Fig. 2a). Topologies 4 and 5 were recovered by both data types, but only topology 5 was robust to both data type and analysis, having been recovered by IQ-TREE analysis of amino acid and codon alignments, and ASTRAL analysis of codon-based gene trees (Table 1; Fig. 2a). Moreover, topology 5 was also recovered by IQ-TREE analysis with the PMSF model, with almost all branches in the *Thalassiosira* grade receiving maximum support (Fig. 1, branches A–E). Notably, the PMSF analysis also recovered a monophyletic *Stephanodiscus*, which matches expectations based on morphology (Theriot et al. 1987). *Stephanodiscus* was paraphyletic in relation to *Cyclostephanos* in 18 of the 20 species trees. Based on all of these results, we chose topology 5 from the PMSF analysis as the reference species tree (Fig. 1).

### Discordance underlies topological uncertainty

Incongruence among genes trees and alignment sites is an important factor impacting phylogenetic reconstruction (Degnan and Rosenberg 2006b; Mallet et al. 2016). We characterized discordance by calculating gene, site, and quartet concordance factors for each branch (Figs. 1 and 2a; Supplementary Figs. S6 and S7). Gene and site concordance factors (gCF and sCF) represent the proportion of genes or sites that are in agreement with a particular branch in the species tree (Minh et al. 2020a). Gene concordance factors range from 0 to 100, and site concordance factors typically range from 33 to 100, with values near 33 indicative of no signal for that branch (Minh et al. 2020a). Quartet concordance factors (QC) provide a likelihood-based estimate of the relative support at each branch for the three possible resolutions of four taxa (Pease et al. 2018). Quartet concordance factors range from −1 to 1, with positive values showing support for the focal branch, negative values supportive for an alternate quartet, and values of zero indicating equal support among the three possible quartets (Pease et al. 2018).

All three concordance factors were high for most branches in the tree, indicating that a majority of genes, sites, and quartets supported those relationships (Supplementary Figs. S6 and S7). Despite having maximum bootstrap and local posterior probability support, however, concordance factors were low for many of the backbone branches (Fig. 1; Supplementary Figs. S6 and S7), including ones within the *Thalassiosira* grade (Fig. 1, nodes A–E) that affected placement of the freshwater cyclostephanoids (Fig. 2). In this part of the tree, concordance was generally low for the backbone branches (gCF = 4–26; sCF = 32–42; QC = −0.04–0.36) (Figs. 1 and 2; Supplementary Figs. S6 and S7). Gene concordance factors were lowest for branches C and D, the only two branches in the species tree with <100% bootstrap support (Fig. 1). Gene concordance factors were only slightly higher for branches A, B, and E (Fig. 1), which had low site concordance (Fig. 2a) and near-zero quartet concordance factors (Supplementary Fig. S7), consistent with expectations for little to no signal. Site concordance factors did not change appreciably with the CDS12 dataset. However, repeating quartet concordance factor calculations with the CDS12 data revealed minor switches in support for branches C (QC_C_ = 0.004 → −0.006) and E (QC_E_ = −0.04 → 0.07) (Supplementary Fig. S7). These patterns of shifting support, though minor, combined with the lowest site concordance factor for branch E (sCF_E_ = 32) suggest that very few sites support the sister relationship of *Thalassiosira* III and cyclostephanoids, despite its recovery in 16 of the 20 inferred species trees (Figs. 1 and 2a).

For some branches on the species tree, there were more discordant than concordant genes and sites (Fig. 1; Supplementary Figs. S6 and S7), suggesting that more genes and sites supported an alternative relationship. This was the case with *Stephanodiscus*, for example, where more genes supported paraphyly than monophyly, though more sites supported monophyly (Supplementary Figs. S6 and S7). Across the tree, support for alternative relationships was neither strong nor consistent among genes or sites. Illustrative of this, discordance in more than one-third of the tree (29 of 83 branches) was due to paraphyly (Supplementary Fig. S6), indicative of widespread lack of signal in gene trees. Taken together, these analyses highlighted extensive gene and site discordance, some well-supported and much of it not, across Thalassiosirales.

There are both biological (e.g., ILS) and technical causes (e.g., gene tree error) of gene discordance, but the proportion attributable to each factor can be difficult to discern (Morales-Briones et al. 2020; Cai et al. 2021). To better assess the importance of gene tree error in our dataset—whether genes with low phylogenetic signal produced inaccurate gene trees—we recalculated gene and site concordance using just the 1588 amino acid orthologs with the highest percentage of parsimony informative sites (top-PI dataset; Table 1). Average gene concordance increased modestly, from 56.6% to 62.1%, but site concordance was unchanged (Supplemental Fig. S8). Gene concordance factors for branches A, B, and E in the *Thalassiosira* grade increased by 5–7% but were largely unchanged for branches C and D (Supplemental Fig. S8). These increases in gene concordance when using the most signal-rich genes suggest that errors in gene tree estimation contributed to the lack of resolution in several critical branches (Chan et al. 2020; Vanderpool et al. 2020). Deeper nodes in the tree may be more prone to technical errors caused by long-branch attraction, poor alignments, or model misspecification, despite our attempts to minimize these during dataset construction (Supplementary File S1). We found slight negative correlations between node age and both gene concordance (*R*^2^ = 0.10, *P* < 0.01) and site concordance (*R*^2^ = 0.26, *P* < 0.001) (Supplementary Fig. S9), which suggests that older branches more likely suffered from saturation due to recurrent substitutions.

### Placement of the freshwater cyclostephanoids

We used two additional tree-based methods to assess the relative support for topologies 1–5 within the *Thalassiosira* grade (Fig. 2a). Like the original species tree inferences, results of AU tests on concatenated alignments for each dataset in Table 1 largely reflected data type (Table 1), with amino acid characters supporting topology 1 (*P* = 0.01, AU test) and the codon and degenerate codon datasets supporting topology 4 (*P* = 0, AU test) (Supplementary Table S4). We then performed gene genealogy interrogation (GGI) to look for secondary signal supporting one or more of the five competing topologies (Arcila et al. 2017). To do this, we performed constrained gene tree searches on the most information-rich (top-PI) orthologs and compared their likelihoods using the AU test. The GGI test assumes monophyly of the tested clades, so using our time-calibrated tree, we converted branch lengths (in millions of years) to coalescent time units using a range of plausible effective population sizes and generation times for diatoms (Supplementary File S1). The estimated stem branch lengths for the five clades in the *Thalassiosira* grade were all >5 coalescent units, suggesting sufficient time to reach monophyly (Rosenberg 2003). After ranking likelihood scores from the AU tests and selecting constraint topologies with the best score (rank 1 trees), no single topology was strongly favored in a majority of constrained gene trees across the three datasets, implying similar levels of support (Fig. 2b–d; Supplementary Table S5). Support from the amino acid dataset was split between topologies 1 and 4, which were recovered by both summary and concatenation methods (Table 1; Fig. 2b; Supplementary Table S5). The most frequent best-fit topology for the codon and degenerate codon datasets corresponded to topology 4 (Fig. 2c, d), which was originally recovered by concatenation only (Table 1).

Gene genealogy interrogation can also be used to explore the effects of gene tree error on summary quartet methods by filtering the input trees for ASTRAL to include only the highest ranking constrained genes (Arcila et al. 2017; Mirarab 2017). For each of the three datasets, we performed two ASTRAL analyses using as input either all the top scoring (rank 1) constrained gene trees (n_AA_ = 1588, n_CDS12_ = 1570, n_DEGEN_ = 1569) or just the subset that had statistical support (*P* < 0.05) above the AU-based rank 2 topology (n_AA_ = 142, n_CDS12_ = 56, n_DEGEN_ = 71). In all six cases, the inferred trees were consistent with topology 5, despite it being best supported (rank 1) in just 13–16% of the constrained gene trees (Fig. 2b–d; Supplementary Table S5). We originally chose topology 5 as the reference species tree (Fig. 1) because it was recovered by both amino acids and codons, ASTRAL and IQ-TREE analysis with the PMSF model, and because it recovered monophyly of *Stephanodiscus*.

Coalescent theory predicts that in severe cases of ILS, short internal branches can produce gene trees that conflict with the species tree more often than they agree, creating a so-called “anomaly zone” (Degnan and Rosenberg 2006a). Our recovery of a species tree topology that is not the most frequent among the GGI gene trees could indicate that the backbone of the *Thalassiosira* grade lies in the anomaly zone. Despite this, polytomy tests in ASTRAL using each dataset rejected the null hypothesis that any of these branches is a polytomy (*P* < 0.05).

### The temporal sequence of marine–freshwater transitions

Divergence time estimates dated the crown Thalassiosirales to the upper Cretaceous, around 113 Ma (95% CI: 96–120 Ma) (Fig. 3a; Supplementary Fig. S10). One of the two main freshwater lineages, *Cyclotella*, originated in the late Cretaceous (95% CI: 66–86 Ma) and the other freshwater lineage, the cyclostephanoids, originated later in the Eocene (95% CI: 36–48 Ma) (Fig. 3a; Supplementary Fig. S10). Radiation of the *Thalassiosira* grade lineages occurred during the Paleocene, from 57 Ma (95% CI: 48–63 Ma) to 73 Ma (95% CI: 61–80 Ma) (Fig. 3a; Supplementary Fig. S10). The overlapping confidence intervals allow for the possibility that these lineages diverged in much more rapid succession than suggested by their mean ages.

**Figure 3.**
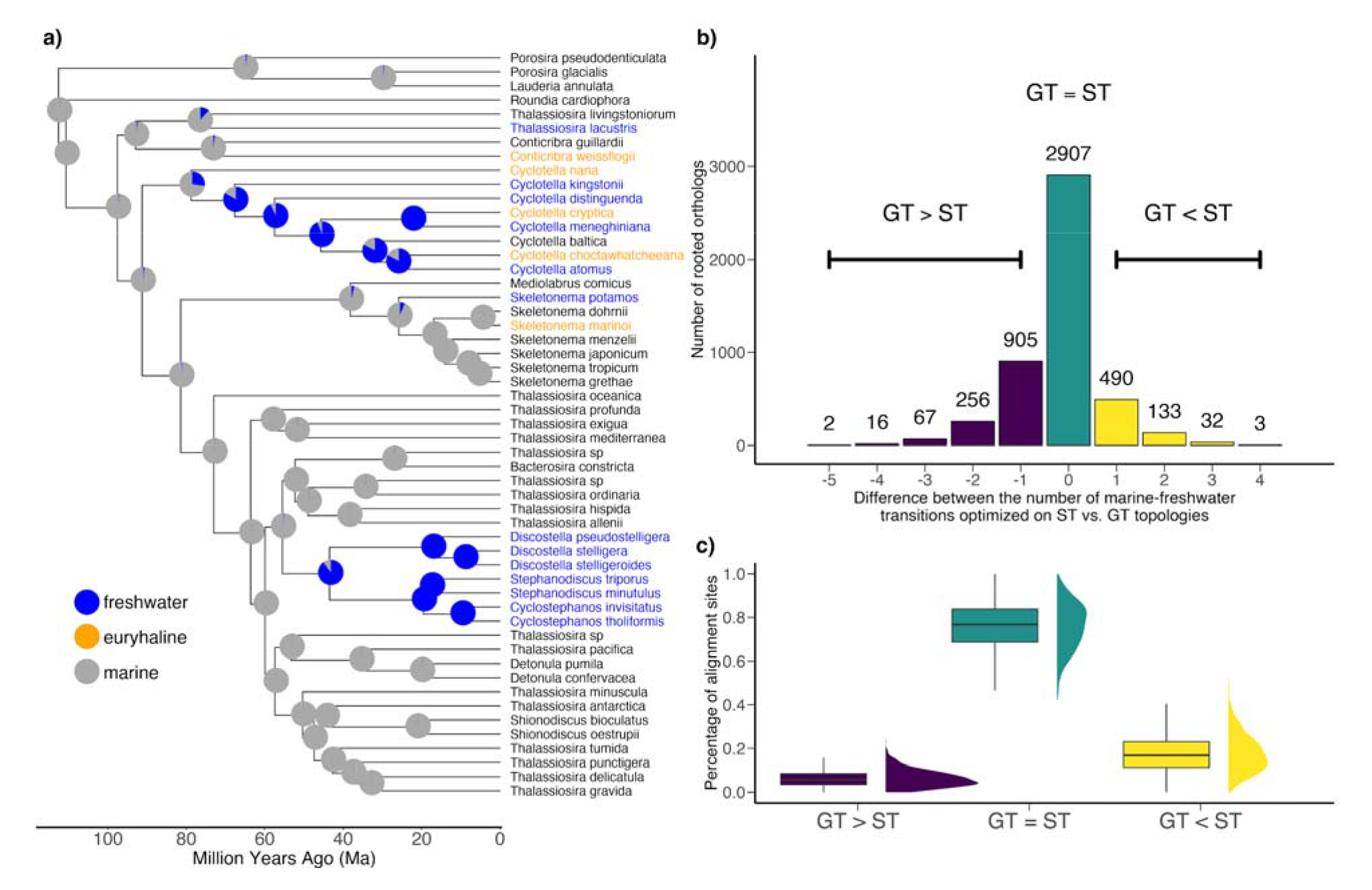
**a)** Divergence times and ancestral state reconstruction of marine and freshwater habitat in Thalassiosirales. Conspecific taxa were removed prior to ancestral state reconstruction, leaving one tip per species. Pie charts denote the probability of each node reconstructed as either marine (grey) or freshwater (blue) using parameters estimated from the HiSSE CID-4 model. Euryhaline taxa were coded as marine for the purposes of ancestral state reconstruction. Divergence times for the full set of taxa can be found in Supplementary Fig. S10. **b)** Summary of the difference between the number of parsimony optimized marine–freshwater transitions on rooted orthologs with the estimated gene tree topology (GT) versus the topology constrained to match the species tree (ST). Bar colors denote whether a GT had greater (purple), equal (green), or fewer (yellow) numbers of transitions relative to the ST. Numbers above bars are the number of rooted orthologs in each bar. **c)** Summary of the percentage of aligned amino acid sites that have greater (purple), equal (green), or fewer (yellow) numbers of state transitions on the GT versus ST.

### Marine–freshwater transitions on gene versus species trees

We used HiSSE to estimate the number of independent marine–freshwater transitions on the time-calibrated species tree (Fig. 3a). The best-fit HiSSE model was a character independent model (CID-4; Supplementary Table S6), which indicates that shifts in diversification rate occurred independently of marine–freshwater transitions. Using parameter estimates from the CID-4 model, we inferred a total of six transitions from marine to freshwaters (Fig. 3a). Given the large number of discordant gene trees (Fig. 1), we next sought to determine whether the reconstructions were impacted by hemiplasy, i.e., marine–freshwater transitions that occurred on branches in a gene tree not shared with the species tree. Narrowly defined, hemiplasy describes discordance due to ILS (Avise and Robinson 2008), but we use the term hemiplasy broadly to describe discordant gene trees with fewer overall habitat transitions than the species tree, regardless of the underlying cause (see Hahn and Nakhleh 2016).

To assess the potential for hemiplasy in our trait reconstructions, we calculated hemiplasy risk factors (HRF), which are the ratio of the probabilities of hemiplasy to homoplasy in different parts of the species tree (Guerrero and Hahn 2018). HRFs were calculated using 12 different sets of coalescent branch lengths that represent a range of scenarios with different effective population size (*N_e_*) and generation times (Supplementary Fig. S11). The risk of hemiplasy ranged from low to nonexistent in nearly all scenarios involving realistic generation times for diatoms (1 day, 7 days, and 1 month), whereas elevated hemiplasy risk was found only under scenarios of unrealistically long generation times (6 months) (Supplementary Fig. S11).

We next assessed whether there was any empirical evidence for hemiplasy in the alignments and gene trees. For each ortholog, we separately optimized, using parsimony, the number of marine–freshwater transitions on the estimated gene tree (GT) and the gene tree constrained to match the species tree topology (ST). We calculated the difference between the number of transitions on the ST versus GT and found a small percentage of orthologs (13.7%) with at least one instance of hemiplasy, broadly defined to include all instances with fewer transitions on the gene versus species tree topology (GT < ST; Fig. 3b). This set of orthologs likely includes topologies affected by hemiplasy due to ILS (Avise and Robinson 2008), homoplasy (convergent or parallel substitutions), and gene tree error. Manual inspection of these gene trees showed that many of them were paraphyletic and occurred between *Thalassiosira lacustris*, *Conticribra weissflogii*, *C. nana*, and the remaining *Cyclotella* taxa, whereas a smaller number were polyphyletic and involved *Cyclotella, Skeletonema potamos,* and the cyclostephanoid clade (Supplemental Table S7). The remaining majority of orthologs either had identical numbers of marine–freshwater transitions (GT = ST), matching the species tree, or had more transitions on the gene tree (GT > ST), which we attribute to error (Fig. 3b).

To determine whether the set of putatively hemiplasious (GT < ST) genes might be related to salinity adaptation, we compared them against a set of genes that were differentially expressed in a low-salinity acclimation experiment using the marine/euryhaline diatom *Skeletonema marinoi* (Pinseel et al. 2022). A total of 532 of the 658 rooted orthologs with GT < ST were also present in the *S. marinoi* genome, and 203 of these (∼38%) were differentially expressed in response to salinity change (Supplementary Table S8). This included genes involved in key pathways and functions important in acclimation to low salinity, such as carotenoid biosynthesis, the xanthophyll cycle, light-harvesting activities, protein chaperones, nitrogen assimilation, regulation of transcription and translation, and the metabolism of polyamines, amino acids, and fatty acids (Supplementary Table S8).

For the same set of 658 orthologs, we then calculated the difference in the number of amino acid transitions that occurred at each alignment site using the inferred GT versus constrained ST topologies (Fig. 3c). Although agnostic to marine–freshwater transitions, this approach measured the impact of gene tree discordance, irrespective of the underlying cause, to overall molecular homoplasy. Using this approach, we found that an average of 18% of amino acid sites within an ortholog had fewer transitions on the GT versus the ST (ST < GT; Fig. 3c).

## DISCUSSION

### Transitions to freshwaters

Marine–freshwater transitions have been key events in the diversification of lineages across the tree of life (Jamy et al. 2022), including diatoms where freshwater taxa have experienced increased rates of both speciation and extinction compared to their marine ancestors (Nakov et al. 2019). Our study focused on a model clade, the Thalassiosirales, which is among the most abundant and diverse lineages in the marine and freshwater plankton and where genetic and genomic resources are readily available (Armbrust et al. 2004; Nawaly et al. 2020; Roberts et al. 2020). Data from the 42 genomes presented here greatly expand the phylogenetic and ecological diversity of sequenced genomes for Thalassiosirales, and diatoms as a whole, and will greatly facilitate efforts to identify the genomic basis of freshwater adaptation in diatoms.

We identified six marine–freshwater transitions in Thalassiosirales and tested whether these transitions were fully independent, owing to separate parallel or convergent mutations (homoplasy) or whether some of the transitions were attributable to hemiplasy, i.e., the sorting of shared ancestral polymorphisms in discordant gene trees (Avise and Robinson 2008). Even in the face of extensive gene tree discordance, the probability of hemiplasy influencing trait reconstructions was virtually nonexistent across the backbone of the tree (Supplementary Fig. S11). The risk of hemiplasy was only elevated under scenarios with unrealistically long generation times for diatoms (Supplementary Fig. S11), many of which can undergo daily cell divisions (Smayda 1969; Tuchman et al. 1984). The large effective population sizes of some microbes, which increase the probability of hemiplasy, may be offset by their rapid generation times, reducing the risk of hemiplasy and highlighting the need for better empirical estimates of *N_e_* and μ in groups like diatoms (Guerrero and Hahn 2018). The relatively few empirical studies of hemiplasy so far have focused on vascular plants or animals with generation times that are years or decades in length (Copetti et al. 2017; Wu et al. 2018; Vizueta et al. 2019). These studies also focused on much younger lineages, making it more likely that shared ancestral polymorphisms could still be detected.

Longer branches subtending separate clades with a shared trait make it more likely that homoplasy, rather than hemiplasy, has occurred (Hahn and Nakhleh 2016). In our case, each of the clades with freshwater taxa have sufficiently long coalescent branch lengths to have achieved monophyly and have had multiple intervening speciation events leading to non-freshwater taxa (Fig. 3a). Other properties of the habitat transitions were also indicative of homoplasy rather than hemiplasy: ancestral state reconstructions were unambiguous, the six freshwater transitions were not paraphyletic, and gene concordance was high on the long internal stem branches subtending the two main freshwater lineages (Hahn and Nakhleh 2016; Wu et al. 2018).

Empirical analyses of gene trees and alignments revealed relatively small numbers of genes and sites with putative hemiplasies (Fig. 3b,c). We identified approximately three-fold fewer amino acid sites with potential hemiplasies than was found in a much younger (10 Ma) radiation of saguaro cacti (Copetti et al. 2017). Given the length of time that has elapsed since the two major freshwater transitions occurred, the probability that shared ancestral polymorphisms persist decreases and the probability that lineage-specific adaptive mutations arise increases (Suh et al. 2015; Zou and Zhang 2015; Mendes et al. 2016). A large number of genes and pathways have been implicated in the response to low salinity (Nakov et al. 2020; Downey et al. 2022; Pinseel et al. 2022), so an adaptive allele in one of the many possible target genes might have made it possible simply to survive in freshwaters initially. In the tens of millions of years since then, any hemiplasious alleles were most likely overwritten in the extant freshwater descendents, leaving a shared ancestral phenotype (e.g., enhanced transport of sodium ions) as the only remaining evidence of the hemiplasy. The divergent evolutionary outcomes also suggest a greater role for homoplasy over hemiplasy. The genus *Cyclotella* includes freshwater, secondarily marine, and generalist euryhaline species that can tolerate a wide range of salinities (Guillard and Ryther 1962; Nakov et al. 2020; Downey et al. 2022). Similarly, most freshwater transitions at the tips of the tree (Fig. 1) involve species with populations that also grow in marine habitats (*Conticribra weissflogii* and *Cyclotella nana*) or can tolerate slightly brackish water (*Thalassiosira lacustris* and *Skeletonema potamos*). The cyclostephanoids are, by contrast, narrowly adapted stenohaline specialists found exclusively in freshwaters, suggestive of a different genetic trajectory into freshwaters.

Although we have treated salinity as a categorical variable, salinity varies along a continuum from freshwater to marine and even hypersaline. Moreover, salinity fluctuations are common in brackish and marine systems, such as coastlines influenced by precipitation and river discharge. Nevertheless, we found that a substantial set of the putatively hemiplasious genes identified here are involved in acclimation to low salinity conditions in one of the species in our analysis, *S. marinoi* (Pinseel et al. 2022), suggesting a possible role for hemiplasy in freshwater adaptation. In addition, adaptation to low or changing salinity may be linked to modifications of gene expression (Bussard et al. 2017; Nakov et al. 2020; Downey et al. 2022; Pinseel et al. 2022), codon usage (Prabha et al. 2017), nucleotide substitution rates (Mitterboeck et al. 2016), transposable element activity (Yuan et al. 2018), epigenetic responses (Artemov et al. 2017), and epistatic interactions (Stern et al. 2022). The genomic resources and phylogenetic framework presented here represent an important advance towards identifying the genes and processes underpinning freshwater adaptation by diatoms.

### The impact of discordance on placement of freshwater clades

Comparative analyses require a strong phylogenetic framework (Felsenstein 1985), so a major goal of this study was to establish a robust phylogenetic hypothesis for Thalassiosirales. Our primarily genome-based analysis of 6262 nuclear orthologs provided better resolution and, superficially, increased support across most of the tree compared to previous analyses based on a few genes (Alverson et al. 2007). Across character types, methods of inference, and different criteria for including characters or taxa, the placement of one of the two principal freshwater clades, *Cyclotella*, was consistent across species trees, with an estimated origin in the late Cretaceous. The placement of the second major freshwater lineage, the cyclostephanoids, was less certain.

The cyclostephanoid clade was placed within a grade of marine species, most of which belong to the polyphyletic genus *Thalassiosira*. These marine *Thalassiosira* were divided among four clades, but the arrangement of these clades and the freshwater cyclostephanoids varied across datasets and analyses. Uncertainty in the backbone relationships for this part of the tree was likely caused by a combination of gene tree error and IL. First, many individual genes contained too little information to confidently resolve deep splits separated by short branch lengths—a finding that is not unique to this dataset (Chan et al. 2020; Arcila et al. 2021). Divergence time estimates suggest that these splits occurred in relatively quick succession, as few as 5 million years. Although many of these bipartitions had consistently weak support, gene concordance factors increased when we analyzed orthologs with the highest phylogenetic signal, implicating gene tree error as a source of instability (Chan et al. 2020; Vanderpool et al. 2020). In other phylogenomic studies impacted by low phylogenetic signal and high gene tree error, gene genealogy interrogation (GGI) has been used to identify majority gene support for a single hypothesis (Hughes et al. 2018; Tea et al. 2021). In our case, no single resolution was supported by a majority of genes, indicative of nodes that are difficult to resolve even with hundreds of genes (Nesi et al. 2021). The best-supported constraint tree identified by GGI differed between amino acid and codon-based gene trees, highlighting conflicting signal even within genes.

Second, in addition to gene tree error, the large number of alternative topologies among gene trees along the *Thalassiosira* grade is also consistent with ILS (Arcila et al. 2017). Gene tree summary methods such as ASTRAL outperform concatenation when ILS is the major source of discordance (Kubatko and Degnan 2007; Roch and Warnow 2015), but summary methods perform poorly when gene tree error is high (Roch and Warnow 2015; Xi et al. 2015). Following Arcila et al. (2017), we attempted to eliminate some of the noise in our dataset by restricting ASTRAL analyses to the top-ranked constrained gene trees and in doing so recovered a backbone topology congruent with one of the few originally recovered by both summary and concatenation methods (topology 5; Table 1 and Fig. 2). Taken together, these results suggest that gene tree error negatively impacted our ASTRAL analyses. After identifying and removing some of that error, we recovered a stronger hypothesis for the placement of freshwater cyclostephanoids within the marine *Thalassiosira* grade.

The anomaly zone describes an especially vexing phylogenetic problem in which short branch lengths are unresolvable, resulting in gene trees that differ from the species more frequently than they agree (Degnan and Rosenberg 2006a). Within the *Thalassiosira* grade, the most common GGI gene tree topologies either did not match the reference species tree or were uninformative for these short branches. This implies that unresolved or weakly supported gene trees are more probable than resolved ones in the *Thalassiosira* grade. Under ILS, branch lengths that exceed the boundaries of the anomaly zone should produce resolved gene trees (Huang and Knowles 2009). Gene tree error like that identified in our dataset can lead to underestimation of coalescent branch lengths by summary methods like ASTRAL (Sayyari and Mirarab 2016; Forthman et al. 2022). These coalescent branch lengths define the anomaly zone boundaries (Degnan and Rosenberg 2006b; Linkem et al. 2016), and underestimates could result in mistaken identification of an anomaly zone where none exists. The *Thalassiosira* grade appears to fall outside of the anomaly zone because our coalescent branch length estimates exceed the theoretical boundaries (Degnan and Rosenberg 2006b).

Factors that are more poorly known in diatoms might also have contributed to the high levels of discordance. We attempted to minimize technical sources of gene tree error during dataset construction (Supplemental File S1), but other sources are more difficult to discern (Cai et al. 2021). These include hybridization and polyploidy, gene duplication and loss, or recombination. Although hybridization does occur in diatoms (Casteleyn et al. 2009; Koester et al. 2010; Tanaka et al. 2015; Çiftçi et al. 2022), it is not especially well studied. There is evidence for an ancient allopolyploidy event early on in the evolutionary history of Thalassiosirales (Parks et al. 2018). Many methods that identify hybridization based on gene trees (Edelman et al. 2019; Vanderpool et al. 2020) or site patterns (Blischak et al. 2018) can be confounded by ancestral population structure, substitutional saturation, or ghost lineages (Slatkin and Pollack 2008; Tricou et al. 2022), all of which are poorly characterized in diatoms. High GC content regions have been linked to higher recombination rates (Kent et al. 2012; Lartillot 2013) and can lead to increased discordance (Pease and Hahn 2013). Despite high levels of variation in GC content across Thalassiosirales, intralocus recombination was detected in a vanishingly small (<0.001%) percent of the 6262 orthologs in our analysis (results available on Zenodo). In addition to showing whether any of these factors have contributed to gene tree discordance, a more thorough exploration of each one will fill important gaps in our understanding of diatom evolution.

### Codon bias and amino acid composition in freshwater diatoms

Thalassiosirales includes divergence times across a timescale ranging from thousands (Theriot et al. 2006) to tens of millions of years ago (Fig. 3a), which led us to explore the utility of both amino acid and nucleotide characters for resolving phylogenetic relationships. Amino acids are less susceptible to saturation and useful for resolving deep relationships (Philippe et al. 2011; Rota-Stabelli et al. 2012), whereas nucleotides contain more information to resolve recent divergences (Simmons et al. 2002; Townsend et al. 2008). Both data types recovered the vast majority of relationships consistently and with strong support, while at the same time revealing similar patterns of discordance along the backbone of the tree. In many cases, however, they differed in their resolutions of the most recalcitrant parts of the tree. Different schemes to mitigate the effects of saturation in the codon datasets also produced disagreements within the *Thalassiosira* grade. Disagreements between character types within the same dataset, such as those within the *Thalassiosira* grade here, have been found in other groups as well (Gillung et al. 2018; Skinner et al. 2020).

Almost every analysis of the amino acid dataset—including species trees, concordance factors, and AU tests—supported the placement of cyclostephanoids as sister to the remaining *Thalassiosira* clades, but nucleotide analyses placed them with *Thalassiosira* III (Fig. 2). Discordance caused by codon usage bias and differences in amino acid composition might account for this discrepancy. An association between codon bias and ecology has been demonstrated in a broad diversity of microbes, where species that share an ecological niche have similar codon usage, independent of phylogeny (Botzman and Margalit 2011; Roller et al. 2013; Arella et al. 2021). Differences in codon usage between marine and freshwater prokaryotes have been described (Cabello-Yeves and Rodriguez-Valera 2019), and we discovered differences in both codon usage and amino acid composition between marine and freshwater diatoms. The amino acid compositions of distantly related freshwater lineages might be sufficiently similar to cause amino acid characters to support the “sister to the rest” placement of cyclostephanoids. Protein sites with different structural, functional, or selective constraints can lead to differences in amino acid composition between species (Villar and Kauvar 1994; Youssef et al. 2021), something that is not accounted for by standard empirical protein models and may have led to artifacts in our gene and species tree inferences (Wang et al. 2018). When we applied the PMSF model, which accounts for compositional heterogeneity in amino acid sites, cyclostephanoids were placed as sister to *Thalassiosira* III, in agreement with the codon datasets. The similarity in codon usage and amino acid composition between distantly related freshwater diatoms merits further study into the causes and functional significance, if any.

## Conclusions

The vast differences between marine and freshwaters result in strong selective pressures on freshwater colonists. Low salinity provokes a broad range of physiological and metabolic responses in diatoms (Nakov et al. 2020; Downey et al. 2022; Pinseel et al. 2022), but the current genetic architectures of freshwater adaptation reflect tens of millions of years of optimization and change since the earliest transitions. As a result, it may be difficult to identify specific alleles—whatever the process that generated them—that currently allow these diatoms to thrive in freshwaters. Nevertheless, the phylogenomic analyses presented here suggest that convergent or parallel substitutions likely played a more important role than hemiplasy in facilitating freshwater colonizations in this group of diatoms. The vast new genomic resources and phylogenetic framework presented here represent an important step forward in addressing these types of questions to better understand how diatoms have made this complex ecological transition appear to be so superficially simple.

## SUPPLEMENTAL MATERIAL

Supplementary Files, Figures, and Tables are available from the Dryad Digital Repository: https://doi.org/10.5061/dryad.7m0cfxpxp. Voucher images, proteomes, alignments, trees, log files, and code have been deposited in Zenodo: https://doi.org/10.5281/zenodo.7713227.

## Supporting information

Supplementary Materials description

## ACKNOWLEDGEMENTS

We thank Ed Theriot for sharing several culture strains, and we thank Matt Ashworth and Jeffery Stone for help with scanning electron microscopy.

