## Supplementary figures and images for "Resolving marine–freshwater transitions by diatoms through a fog of discordant gene trees"

### Supplementary Fig. S1

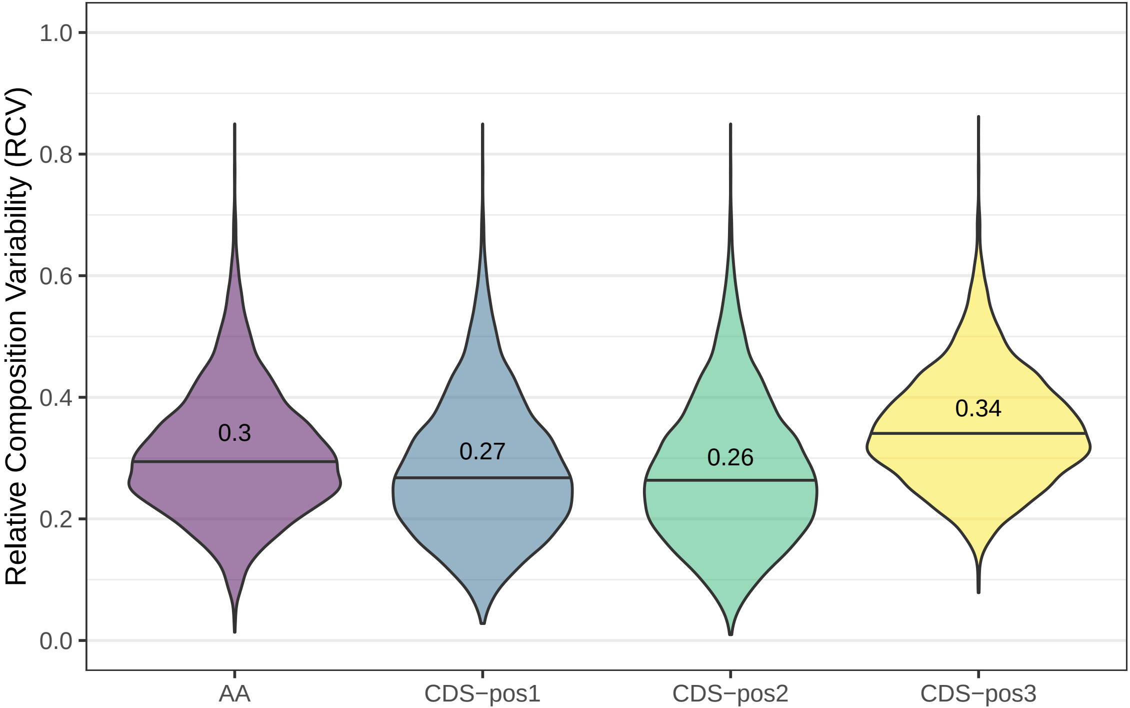

### Supplementary Fig. S2

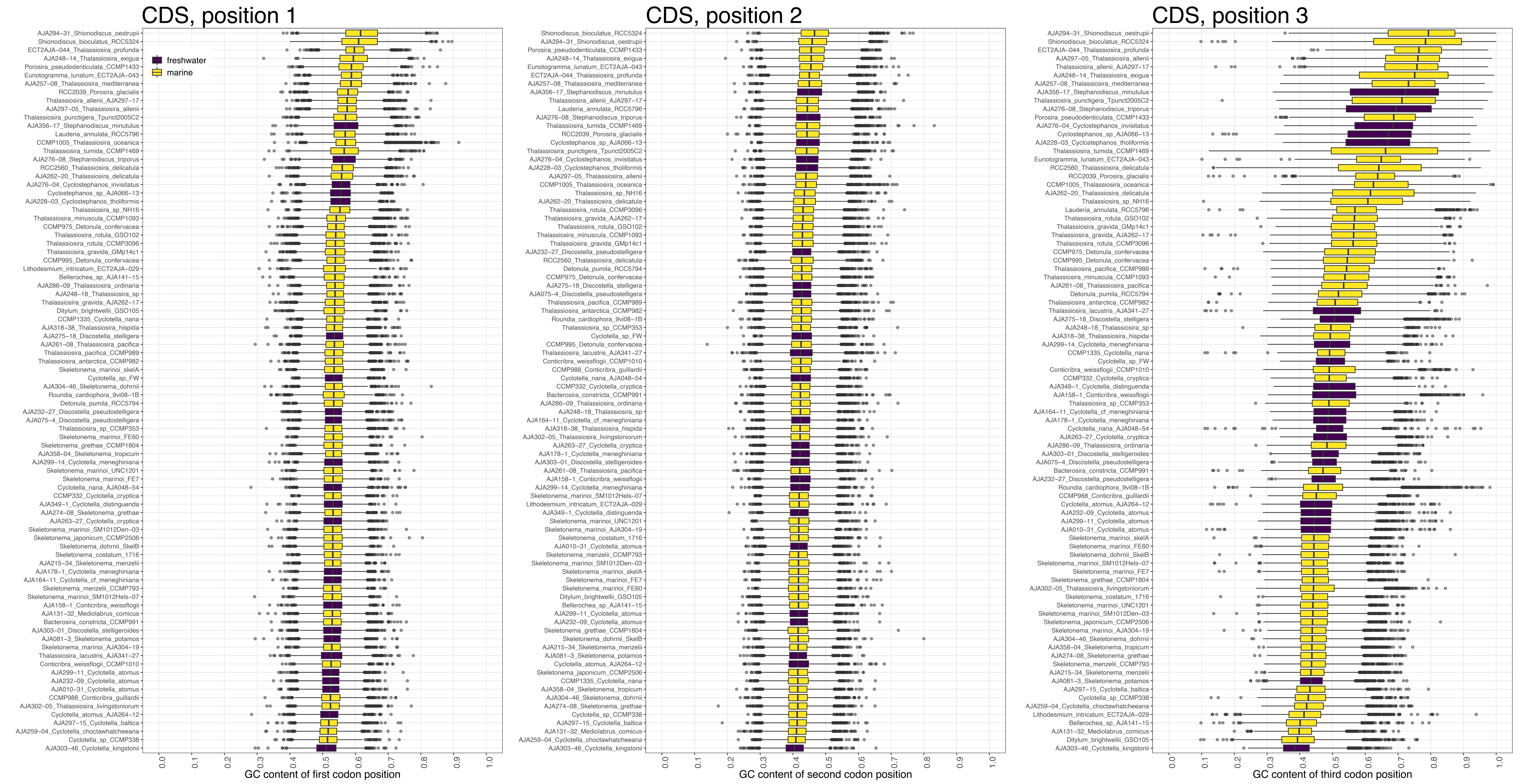

### Supplementary Fig. S3

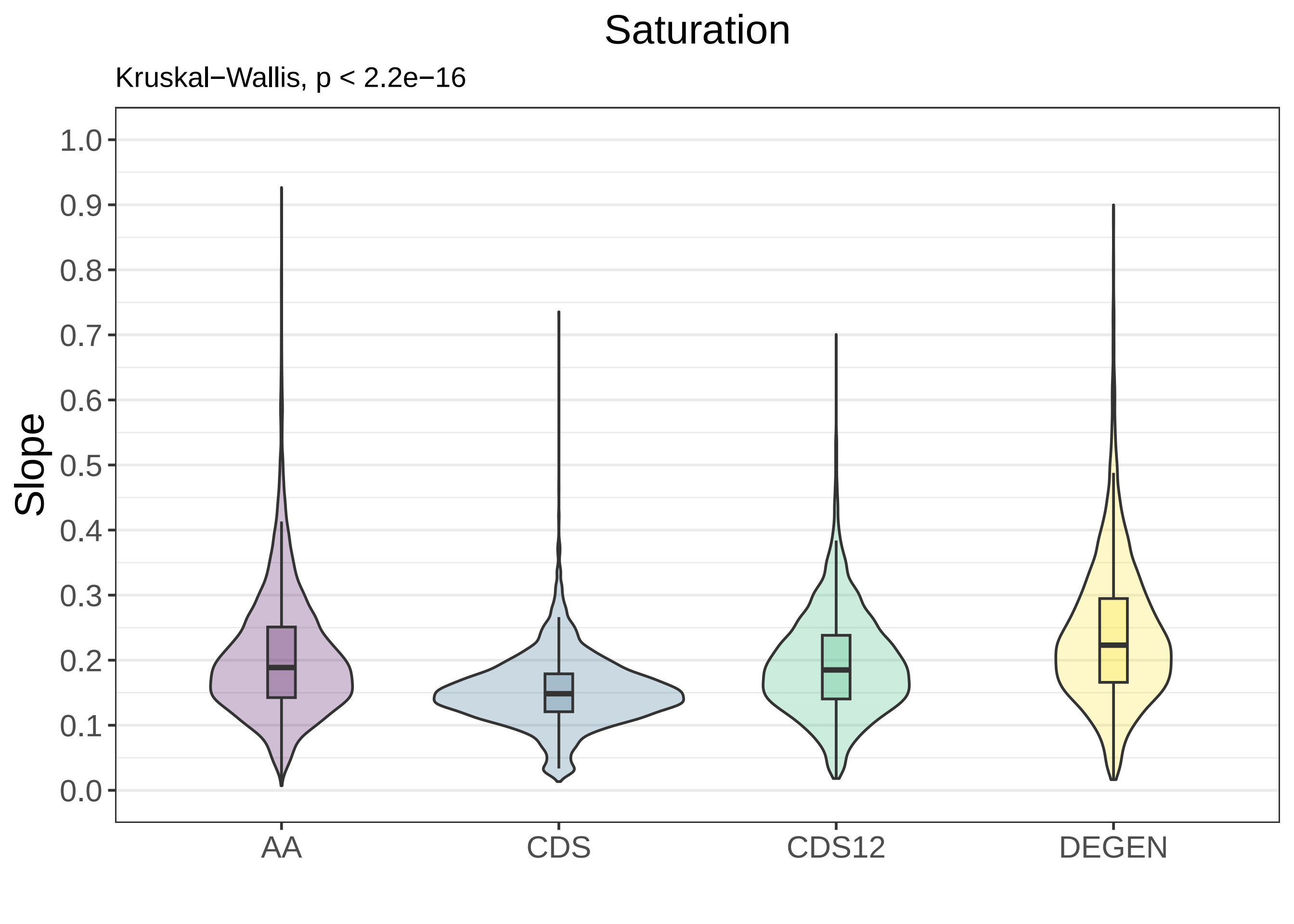

### Supplementary Fig. S4

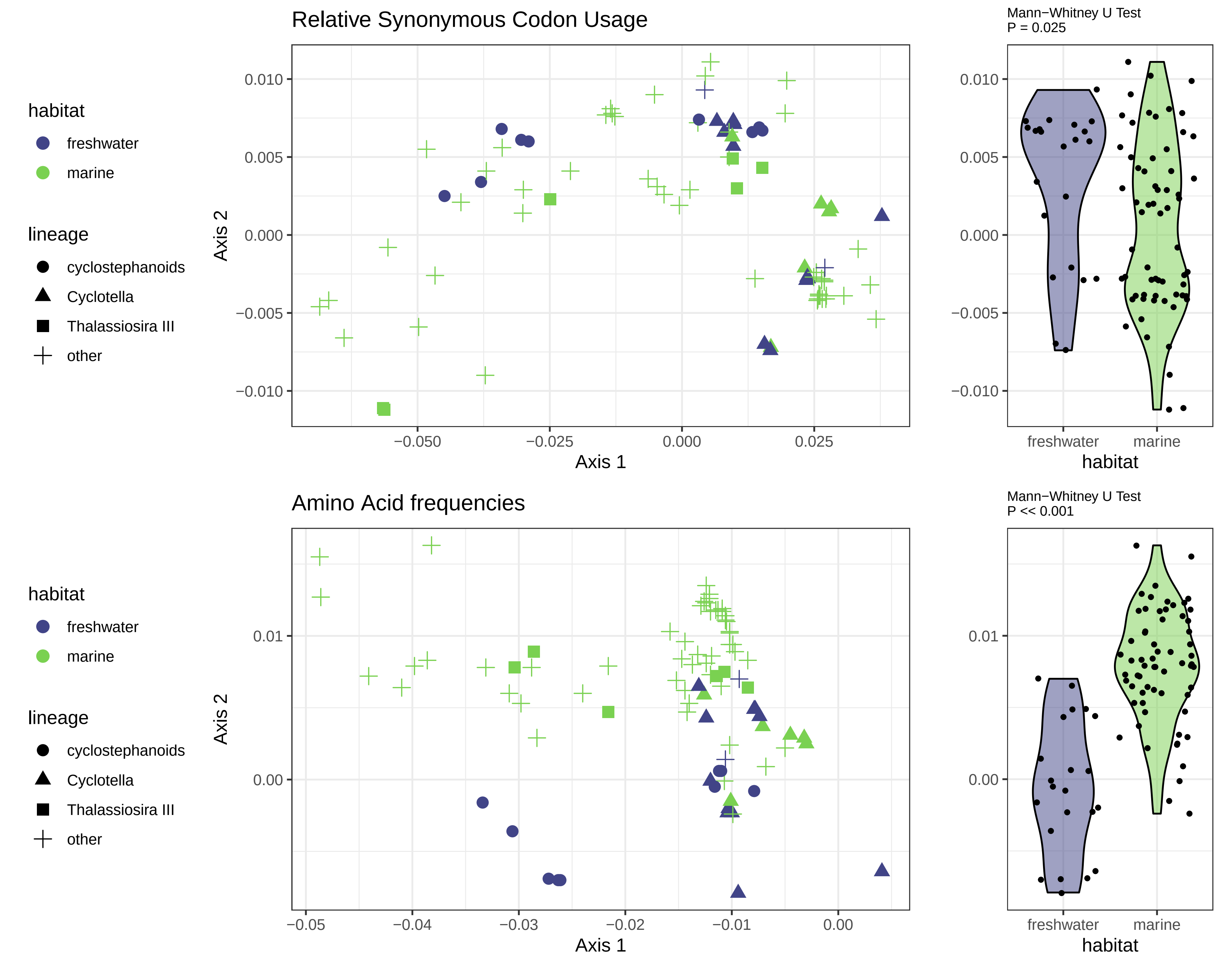

### Supplementary Fig. S5

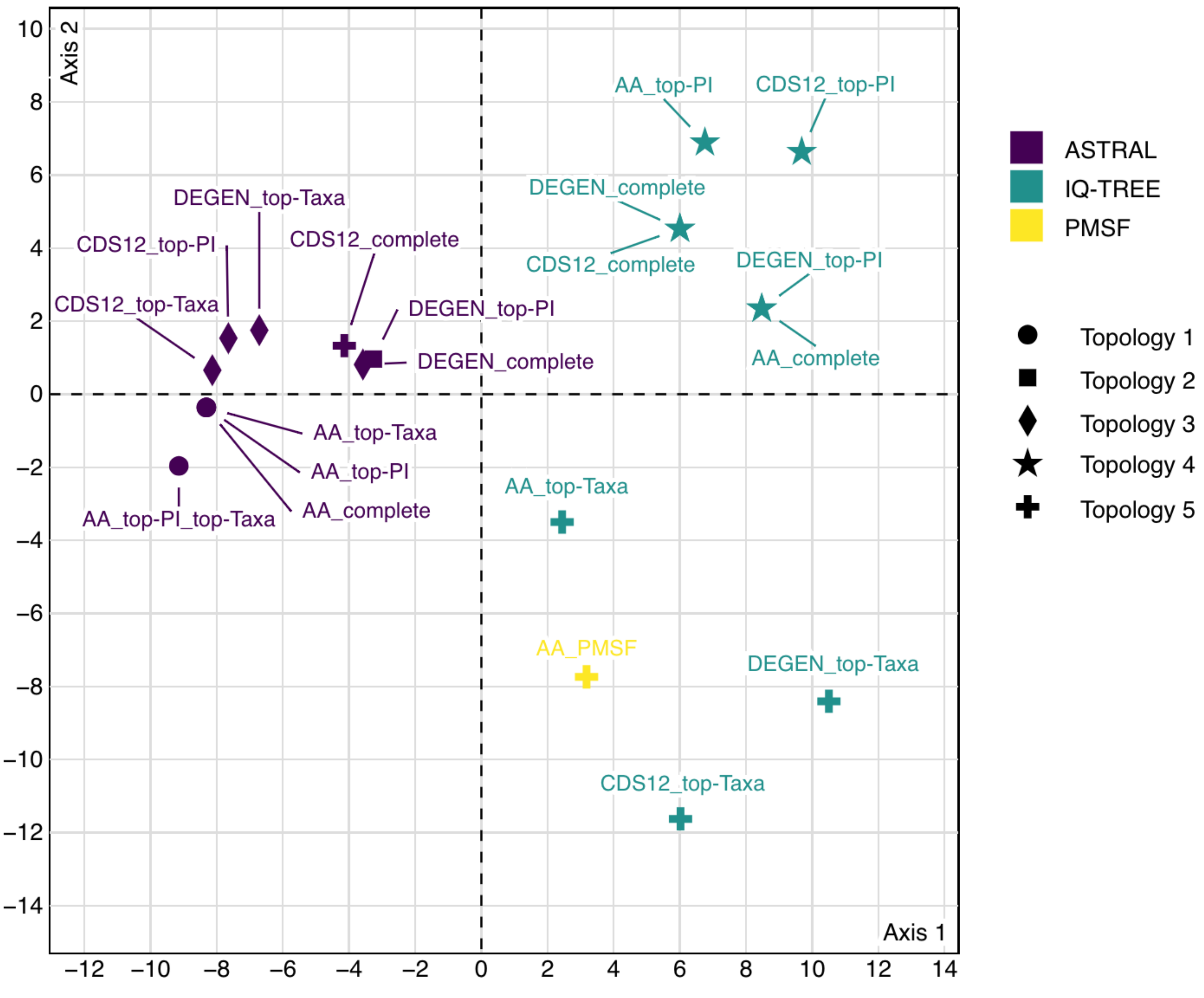

### Supplementary Fig. S6

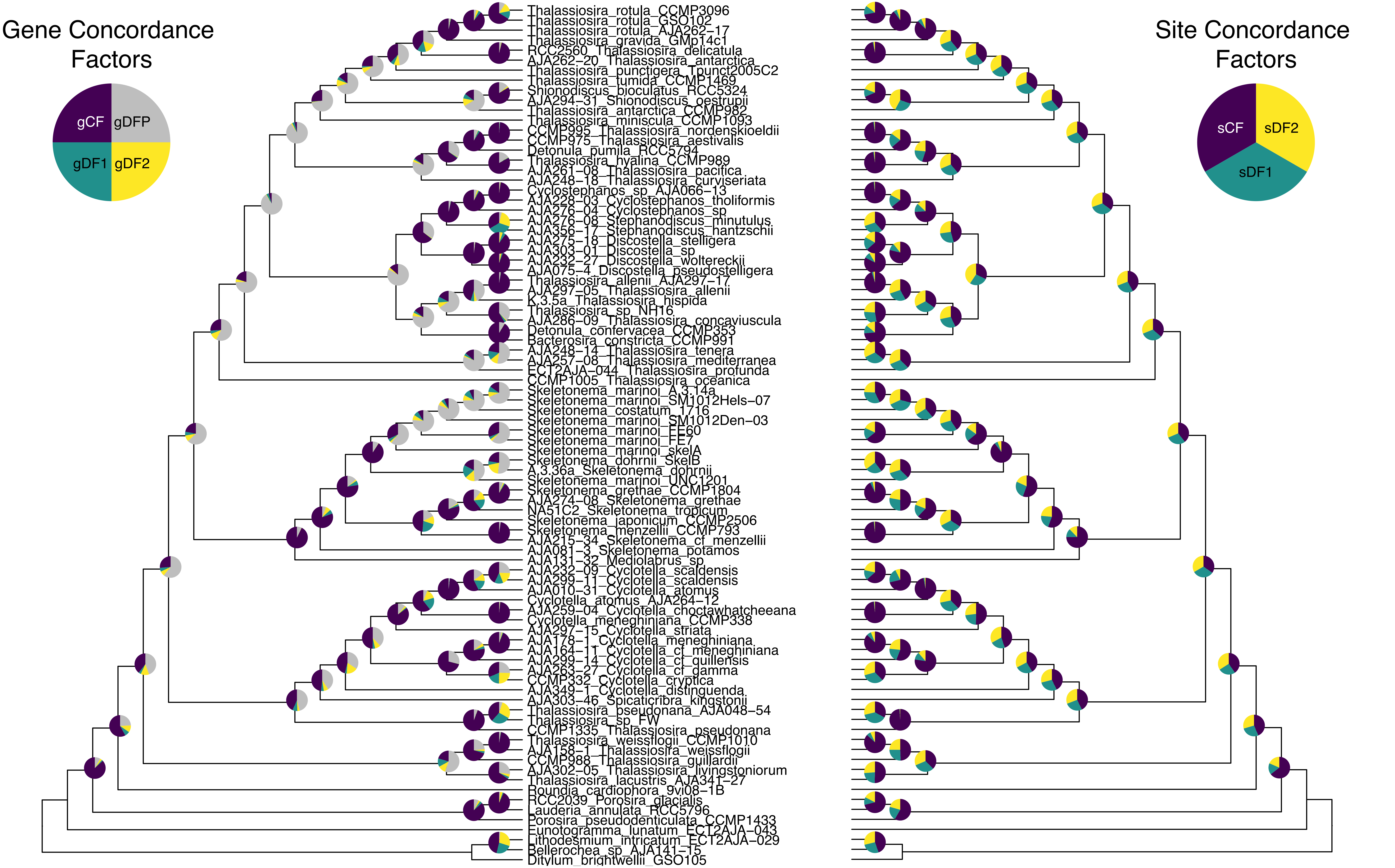

### Supplementary Fig. S7

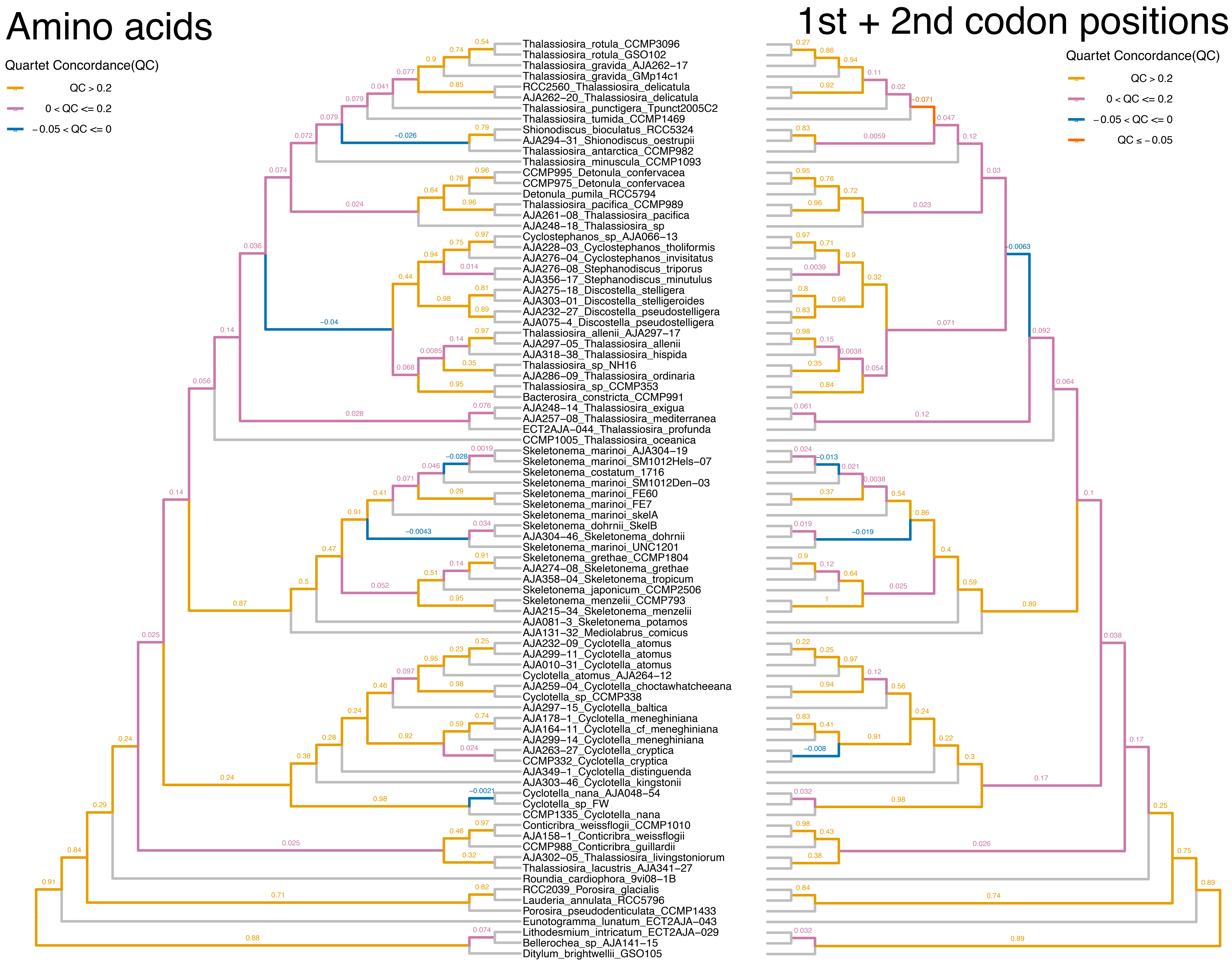

### Supplementary Fig. S8

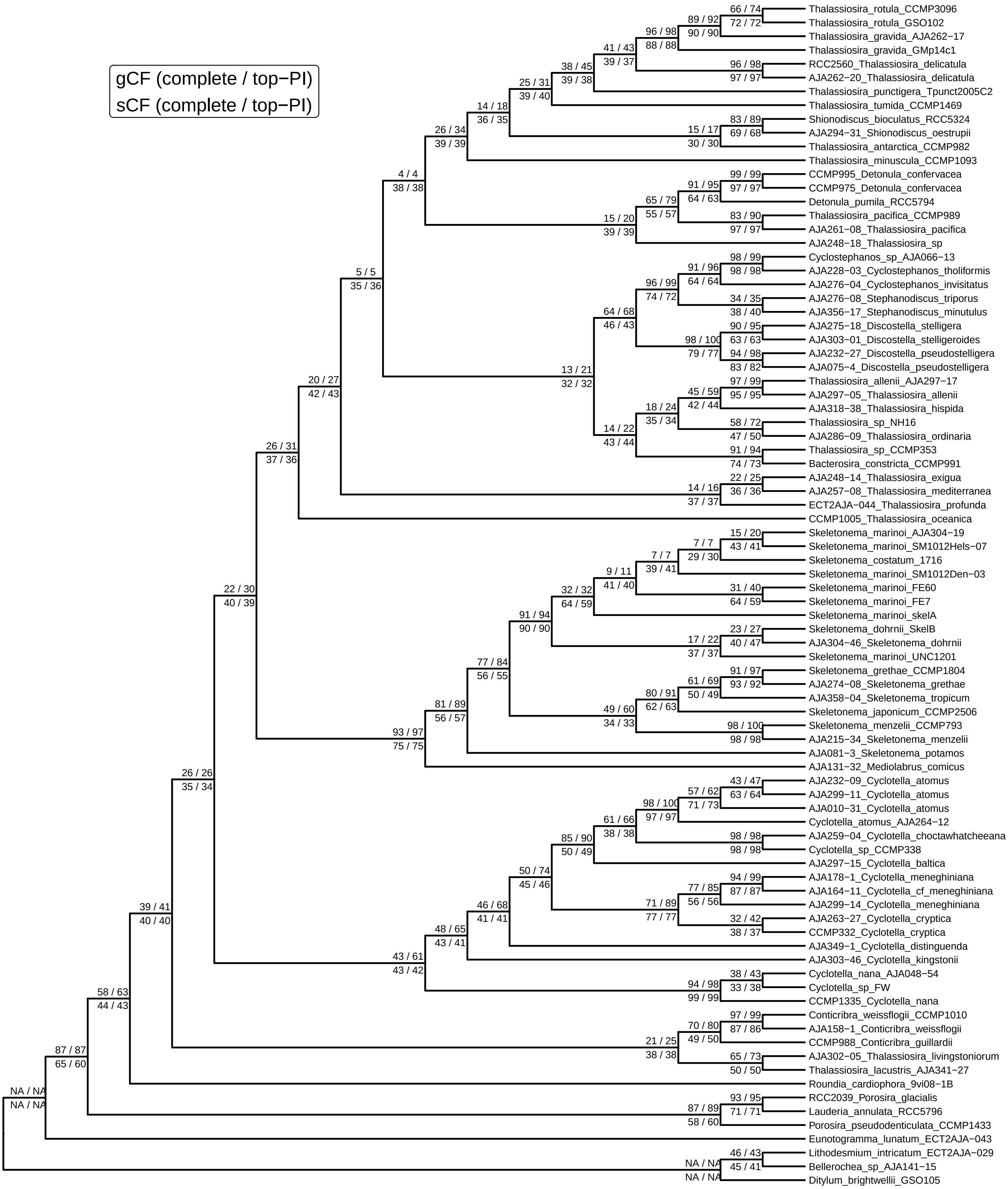

### Supplementary Fig. S9

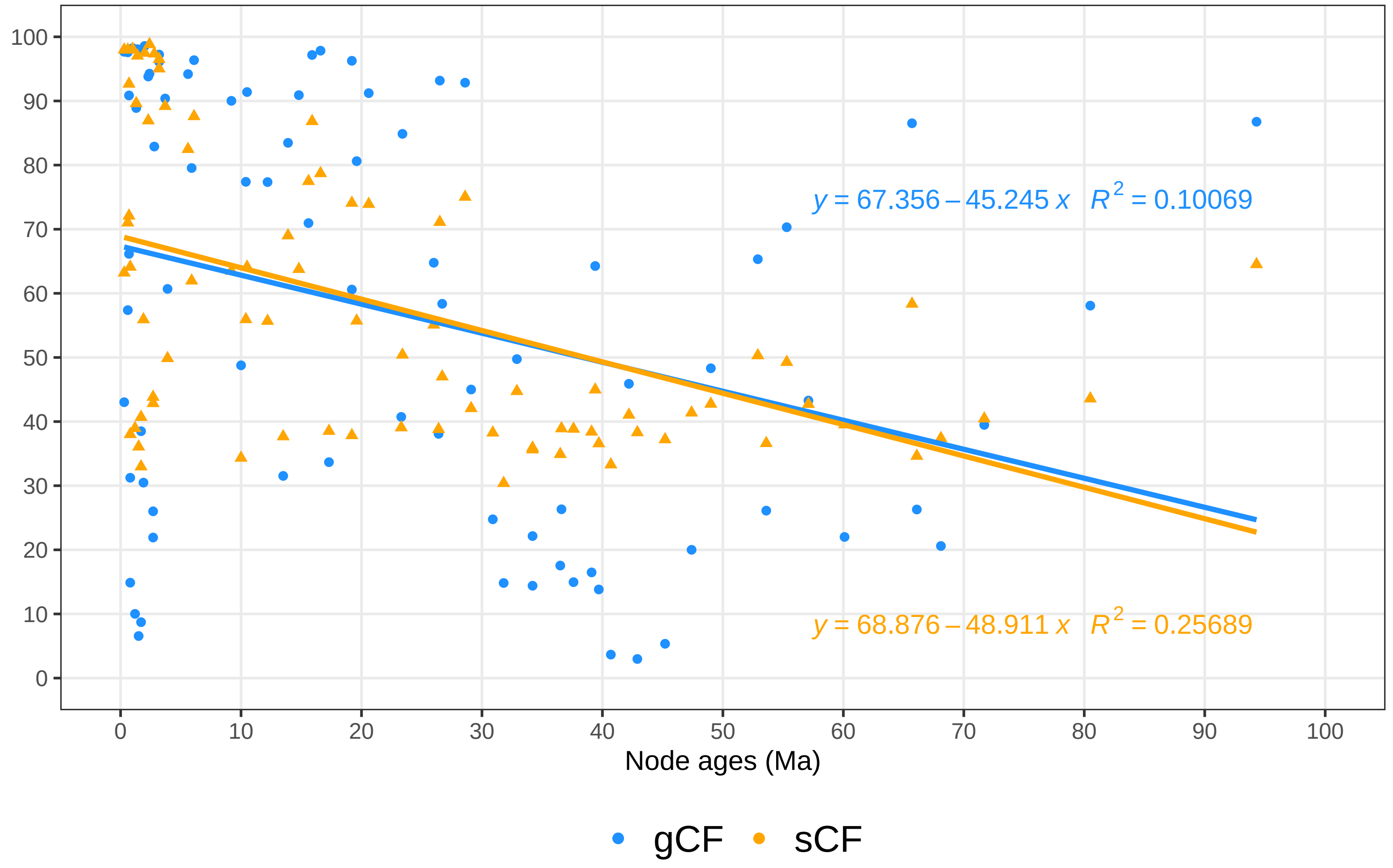

### Supplementary Fig. S10

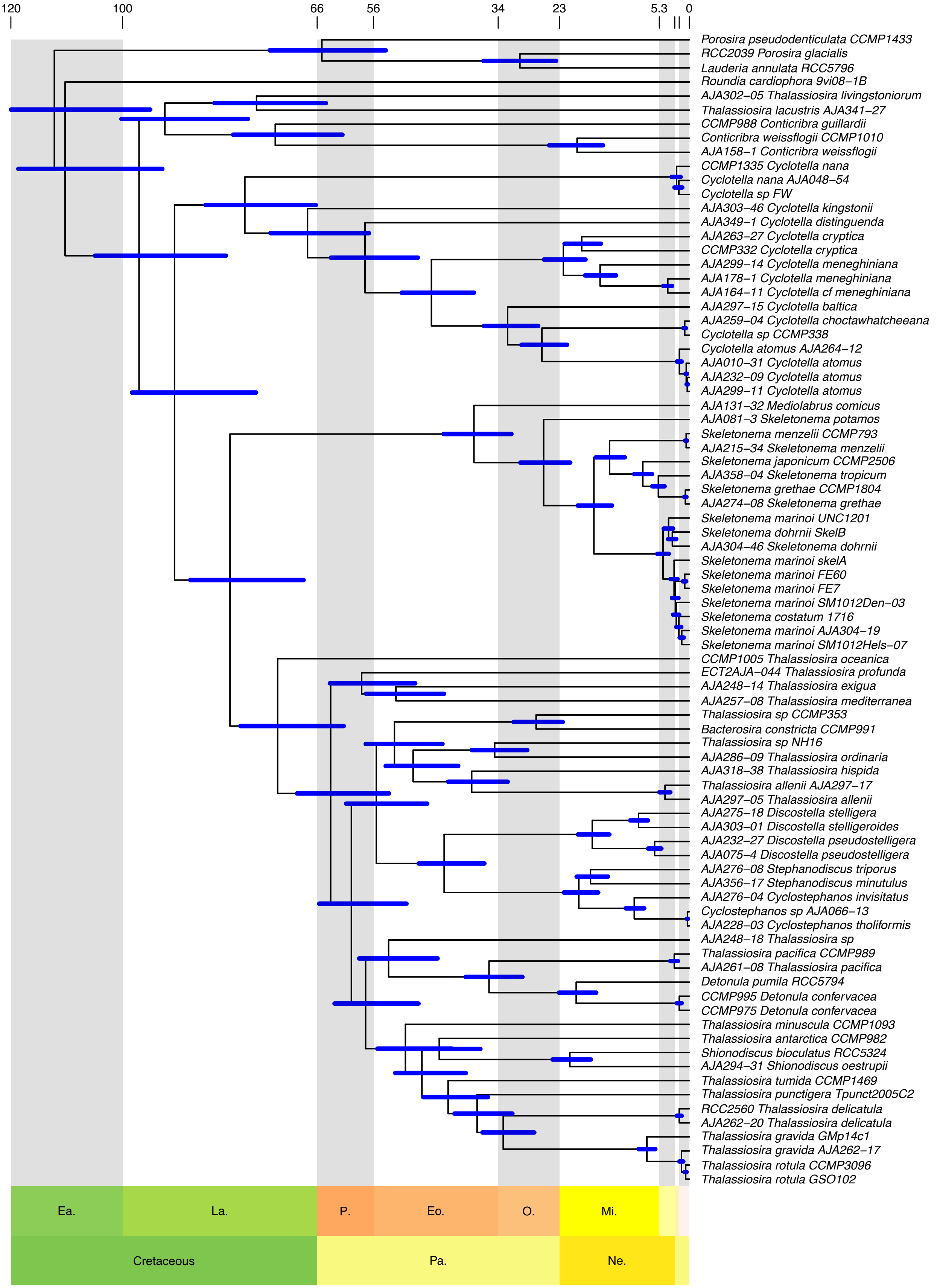

### Supplementary Fig. S11

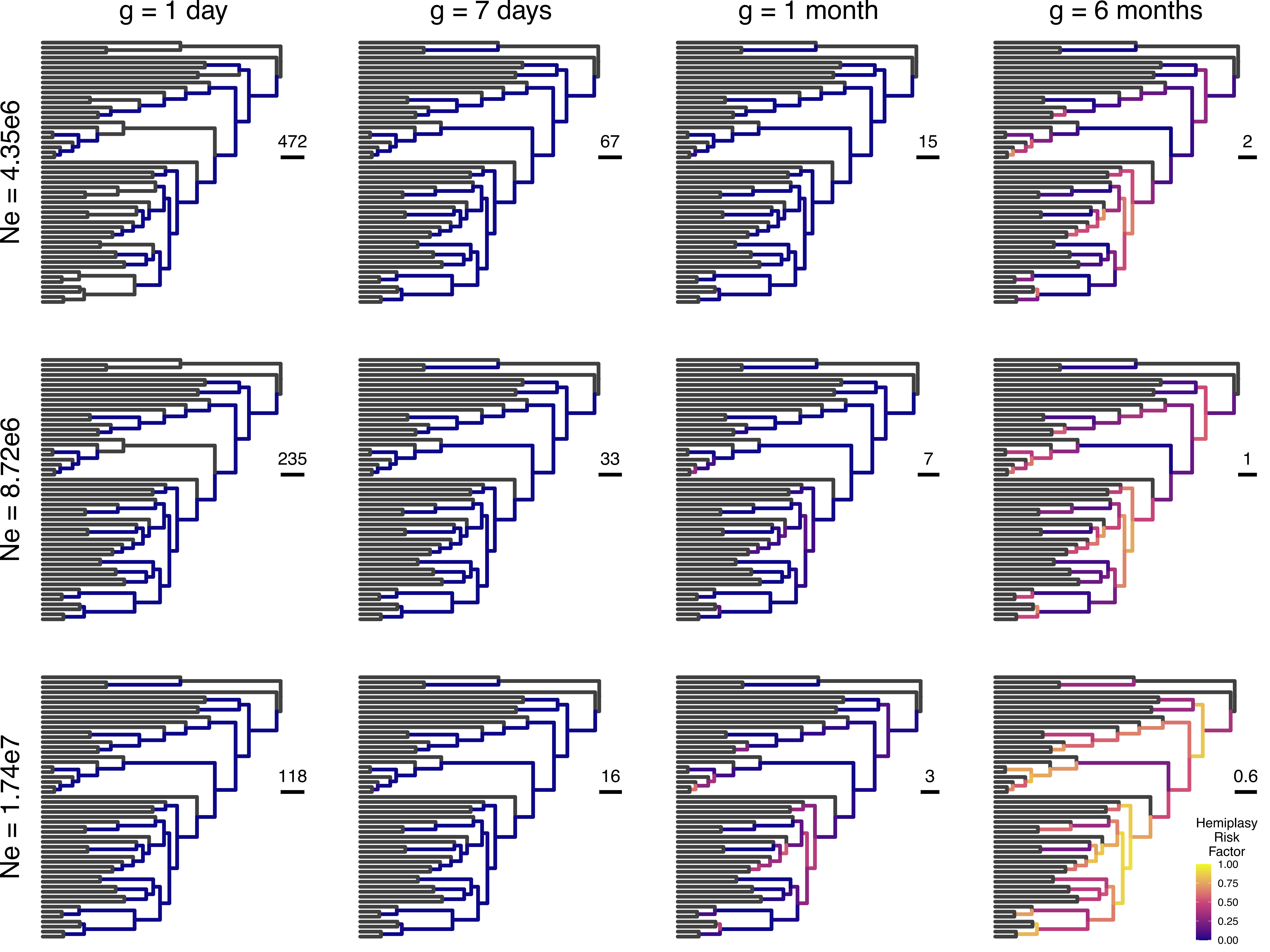
