## Supplementary File S1 for "Resolving marine–freshwater transitions by diatoms through a fog of discordant gene trees"

### SUPPLEMENTARY FILE S1: MATERIALS AND METHODS

#### *Strain Collection and Culturing*

Eight strains were acquired from the National Center for Marine Algae and Microbiota (NCMA) in the United States. Five strains were acquired from the Roscoff Culture Collection (RCC) in France. Additional strains were collected from a variety of locales in the United States and isolated into monoclonal cultures. Marine strains were grown in L1 medium (Guillard and Hargraves 1993) and freshwater strains were grown in WC medium (Guillard and Lorenzen 1972). Cells were grown in batch culture at varying temperatures from 4-21 ℃ on a 12-12 hour light-dark cycle. Collection data, growth conditions, and taxonomic identification for all cultures are provided in Supplementary Table S1. Voucher images are provided on Dryad. Cultures newly isolated in the Alverson Lab have been submitted to the public culture collections at NCMA.

#### *Transcriptome Sequencing and Assembly*

We extracted total RNA using the RNeasy Plant Mini Kit (Qiagen) from strains growing at log-phase growth. We assessed RNA quantity using the Qubit (RNA BR Assay kit; Thermo Fisher) and prepared mRNA libraries using either the TruSeq RNA Kit (Illumina), KAPA stranded mRNA-Seq Kit (Roche), or the KAPA HyperPlus Kit (Roche). We barcoded each library with single or dual indices and assessed their quality using the TapeStation (Agilent).

Library pools were sequenced on an Illumina HiSeq4000 at the Beijing Genomics Institute (BGI) or an Illumina NovaSeq at the University of Chicago Genomics Facility.

We corrected sequencing errors in the raw RNA-seq reads using Rcorrector v.1.0.1 (Song and Florea 2015) with a k-mer size of 31. We trimmed and quality filtered the corrected reads using Trimmomatic v.0.36 (Bolger et al. 2014) with options 'ILLUMINACLIP:TruSeq3-PE-2.fa:2:40:15 LEADING:2 TRAILING:2 SLIDINGWINDOW:4:2 MINLEN:30'. We then filtered reads belonging to vector contaminants, ribosomal RNA, and organelles using Bowtie2 v.2.2.3 (Langmead and Salzberg 2012; Langmead et al. 2019) against custom databases. We assembled the clean reads into transcripts using Trinity v.2.10.0 (Grabherr et al. 2011; Haas et al. 2013), under default options. We assessed transcript quality using TransRate v.1.0.3 (Smith-Unna et al. 2016) and filtered low-supported transcripts (sCnuc <0.25, sCcov <0.25, sCord <0.5). We removed chimeric transcripts using a BLAST-based approach (Yang and Smith 2013). We next clustered the transcripts into putative genes using Corset v.1.0.9 (Davidson and Oshlack 2014), keeping the longest transcript for each cluster as the representative. We then predicted open reading frames using TransDecoder v.5.5.0 (https://github.com/TransDecoder/TransDecoder), while providing homology evidence from a Diamond BLASTp v.2.0.1 (Buchfink et al. 2015) search against the SwissProt database (release 2021_01; (UniProt Consortium 2021). The single-best ORF per transcript was kept ('--single_best_only') and translated into amino acids. Finally, we removed contaminant sequences identified by CroCo v.1.1 (Simion et al. 2018).

We additionally downloaded the reads for the Thalassiosirales strains included in the Marine Microbial Eukaryote Transcriptome Sequencing Project (Keeling et al. 2014) from NCBI BioProject PRJNA231566. We corrected, trimmed, and assembled the reads as above.

*Genome Sequencing, Assembly, and Gene Prediction*

We extracted total genomic DNA using either a modified CTAB protocol (Doyle and Doyle 1987) or a Qiagen DNeasy Plant Kit (Qiagen), followed by RNase treatment. We assessed DNA quality and quantity using the Nanodrop (Thermo Fisher) and Qubit (dsDNA HS Assay kit; Thermo Fisher). We prepared libraries from DNA using the KAPA HyperPrep Kit (Roche), barcoded each with dual indices, and assessed their quality using the TapeStation (Agilent). Library pools were sequenced on either an Illumina HiSeq4000 or NovaSeq at the University of Chicago Genomics Facility.

We quality trimmed and quality filtered the raw reads using Trimmomatic with options 'ILLUMINACLIP:TruSeq3-PE-2.fa:2:30:10 LEADING:3 TRAILING:3 SLIDINGWINDOW:4:15 MINLEN:50'. We then corrected the filtered reads for sequencing errors with BayesHammer (Nikolenko et al. 2013) and assembled them with SPAdes v.3.12.0 (Bankevich et al. 2012), using the default k-mer lengths of 21, 33, and 55. We visualized and removed contaminants from the resulting assemblies using the Blobtools v.1.1 pipeline (Laetsch and Blaxter 2017). Read coverage was estimated by mapping the corrected reads against the assembly using minimap2 v.2.10-r761 (Li 2018). We assigned a taxonomic identity to each contig using a Diamond BLASTx search against the UniProt Reference Proteomes database (releases 2019_06 or 2020_04) using options '--max-target-seqs 1 --sensitive --evalue 1e-25 --outfmt 6'. We filtered each assembly to remove: (1) small (<500 bp) contigs with no taxonomic hit, and (2) contigs with taxonomic assignment to bacteria, archaea, or viruses. We then extracted the strictly paired reads from the kept contigs (non-contaminant) and reassembled them with SPAdes. We again visualized the new assemblies with Blobtools and additional contaminant contigs were removed using the same procedure. We then created scaffolds from each assembly with Rascaf v.1.0.2 (Song et al. 2016), using RNA-seq reads, followed by SSPACE v.3.0 (Boetzer et al. 2011), using DNA-seq reads. Lastly, we performed contig extension and gap closure using GapCloser v.1.12 (Luo et al. 2012) with both the RNA- and DNA-seq reads. We assessed each SPAdes assembly for quality and completeness using QUAST v.5.0.0 (Gurevich et al. 2013) and BUSCO v.5.1.3 (genome mode; stramenopiles_odb10 dataset; (Simão et al. 2015). To identify chloroplast and mitochondrial contigs, we used NCBI BLASTn v.2.12.0+ (Camacho et al. 2009) to search each assembly against a set of Thalassiosirales chloroplast and mitochondrial genomes downloaded from GenBank (Accessions: NC_025314.1, NC_025312.1, NC_014808.1, NC_008589.1, NC_038005.1, NC_007405.1, NC_028615.1). We subsequently removed contigs with organellar BLAST hits.

We predicted gene models for each genome using the MAKER2 pipeline v.2.31.10 (Cantarel et al. 2008; Holt and Yandell 2011). For evidence during the first round of MAKER, we used the assembled transcriptome coding sequences from the genome of interest (est2genome) and protein homologs from other sequenced diatom genomes (protein2genome; proteomes from *Cyclotella nana, Phaeodactylum tricornutum, Fragilariopsis cylindrus,* and *Thalassiosira oceanica*; downloaded from the Joint Genomes Institute [JGI] PhycoCosm resource; (Grigoriev et al. 2021). For some genomes without corresponding RNA-seq evidence, we used the transcriptome from an alternate strain as evidence. For the second round of MAKER, we used the CDS and protein alignments from the first round to train SNAP v.2006-07-28 (Korf 2004) models for *ab initio* gene prediction. We also included the trained Augustus (Stanke et al. 2008) models developed for *Cyclotella cryptica* CCMP332 (Roberts et al. 2020) as a second *ab initio* gene predictor. For the third round of MAKER, we retrained

SNAP using the gene prediction results from the second round. We assessed each round of MAKER using BUSCO (protein mode; stramenopiles_odb10 dataset) and the count of predicted genes with Annotation Edit Distance (AED) <0.5 (Holt and Yandell 2011). We considered either the second or third MAKER rounds the best if it had the highest number of recovered BUSCO genes and highest number of gene models with AED <0.5. We generated the transcript and translated protein sequences for each gene model using MAKER tools. We also generated files containing the coding sequences for the gene models using GffRead (Pertea and Pertea 2020).

We downloaded genomes and gene models for the previously assembled Thalassiosirales genomes *Cyclotella nana* CCMP1335, *Thalassiosira oceanica* CCMP1005, and *Cyclotella cryptica* CCMP332 (Armbrust et al. 2004; Lommer et al. 2012; Roberts et al. 2020). We downloaded the genome and gene models for CCMP1335 and CCMP1005 from the JGI PhycoCosm resource. We included a reduced version of the CCMP1005 genome and gene models in which redundant contigs and gene models were removed (available from DOI: 10.5281/zenodo.4589594). For all proteomes and coding sequences derived from the newly assembled genomes in this study, we removed all alternative isoforms and kept only the longest protein from each genome locus using the R Bioconductor package *Biostrings* v.2.62.0 (DOI:10.18129/B9.bioc.Biostrings).

*Multiple Sequence Alignment and Gene Tree Estimation*

Our full dataset contained 44 genomes and 43 transcriptomes, including 82 ingroup and 5 outgroup taxa. Based on Nakov et al. (2018), transcriptomes for the following outgroup taxa were included: *Coscinodiscus* sp. AJA212-04, *Ditylum brightwellii* GSO105, *Bellerochea* sp. AJA141-15, *Lithodesmium undulatum* ECT2AJA-029, and *Eunotogramma intricatum*

ECT2AJA-043. We clustered the predicted proteomes into orthogroups with OrthoFinder v.2.4.0 (Emms and Kelly 2019), under default options. We retained 11,687 orthogroups that contained at least 20% of the ingroup taxa and aligned the amino acid sequences for these orthogroups using MAFFT v.7.304b (Katoh and Standley 2013). For orthogroups with <200 sequences, we used the E-INS-i ('--genafpair --maxiterate 1000') algorithm, otherwise the '--auto' option was used. Aggressive alignment filtering can negatively impact downstream gene tree and species tree estimation by removing potentially informative sites (Tan et al. 2015; Portik and Wiens 2021). We therefore performed light alignment trimming to remove poorly aligned regions with >90% missing data using TrimAl v.1.4 (Capella-Gutiérrez et al. 2009). We then estimated phylogenetic trees for each trimmed alignment using IQ-TREE v.1.6.12 (Minh et al. 2020b) or FastTree v.2.1.10 (Price et al. 2010). For the trimmed alignments containing <1000 sequences, a tree was estimated using IQ-TREE and the best fitting substitution model was determined using ModelFinder (Kalyaanamoorthy et al. 2017). For the trimmed alignments containing >1000 sequences, a tree was estimated with FastTree using the WAG substitution model ('-wag') and rescaling branch lengths using a Gamma likelihood ('-gamma').

We then trimmed and filtered the resulting trees using the scripts from (Yang and Smith 2014) (https://bitbucket.org/yanglab/phylogenomic_dataset_construction/). First, we trimmed long and spurious tips from each tree using TreeShrink v.1.3.7 (Mai and Mirarab 2018) with the 'per-gene' mode. We tested four different quantile values (0.05, 0.01, 0.005, and 0.001) and assessed how many outgroup taxa were removed. To balance the removal of spurious tips with the number of outgroups remaining, we chose a quantile cutoff of 0.01. Next, we masked both mono- and paraphyletic tips that belonged to the same taxon. Finally, after visual inspection of trees, we cut deep paralogs out from the masked trees using a branch length cutoff of 0.5 and requiring a minimum of 20 taxa. We created new sequence files from these trimmed and filtered trees and repeated all steps above for a second time using all of the same options.

After the two rounds of tree trimming and filtering, the resulting trees are considered to be homolog trees, containing both orthologs and paralogs. We aligned the final amino acid homologs using MAFFT and lightly trimmed the alignments as above. For homologs with <500 sequences, we estimated final homolog trees using IQ-TREE, using ModelFinder to determine the best-fit model ('-m TEST'), performing 1000 ultrafast bootstraps to assess branch support ('-bb 1000'), and performing 5 independent tree searches ('--runs 5'). For homologs with >500 sequences, we estimated final homolog trees using FastTree, using the WAG substitution model ('-wag'), performed four rounds of minimum evolution SPR moves ('-spr 4') and a more exhaustive NNI search ('-mlacc 2 -slownni'), and rescaled branch lengths using Gamma likelihood ('-gamma').

To prune out rooted orthologs from the homolog trees, we used the Rooted Ingroup (RT) method of Yang and Smith (Yang and Smith 2014), using *Coscinodiscus* sp. AJA212-04 as the outgroup and keeping orthologs with a minimum of 22 ingroup taxa. For the purposes of the RT algorithm, outgroup taxa *Bellerochea* sp. AJA141-15, *Ditylum brightwellii* GSO105, *Eunotogramma lunatum* ECT2AJA-043, and *Lithodesmium intricatum* ECT2AJA-029 were considered as ingroup taxa. We created new sequence files from the pruned orthologs for both the amino acid and coding sequence data. We then aligned the amino acid sequences using MAFFT under the L-INS-i ('--localpair --maxiterate 1000') algorithm. To create the coding sequence alignments, we took the amino acid alignment and corresponding coding sequences and used PAL2NAL v.14 (Suyama et al. 2006) to create the codon alignments. Consistency errors were found between amino acid and coding sequences for *C. nana* CCMP1335. We manually checked these errors with GeneWise (https://www.ebi.ac.uk/Tools/psa/genewise/; (Birney et al. 2004) and fixed them before rerunning the codon alignments with PAL2NAL. We again lightly trimmed both amino acid and coding sequence alignments to remove columns with >90% missing data. For the trimmed amino acid and coding sequence alignments, we estimated final ortholog trees using IQ-TREE, using ModelFinder to determine the best-fit model for each alignment ('-m TEST'), performing 5 independent tree searches ('--runs 5'), and assessed branch support with 1000 ultrafast bootstraps ('-bb 1000'). We partitioned the coding sequence alignments by codon position. We collapsed gene tree branches with very low support (<33% ultrafast bootstrap support) as this has been demonstrated to improve downstream species tree accuracy (Zhang et al. 2018; Shen et al. 2020).

*Ortholog Summaries*

For the orthologs in each dataset (amino acids and coding sequences), we used AMAS (Borowiec 2016) to calculate the following summary statistics: number of taxa, length of the trimmed alignment, percentage of parsimony-informative (PI) sites, percentage of missing data, and the guanine-cytosine (GC) content for each codon position per taxon per gene. We calculated the average bootstrap branch support and saturation for each ortholog tree using custom R code (R Core Team 2021). We additionally calculated the Relative Composition Variability (RCV; (Phillips and Penny 2003) for each dataset using PhyKit v.1.2.1 (Steenwyk et al. 2021). RCV is the average variability in composition between taxa in an alignment and can be useful for evaluating potential sequence composition bias. Following calculation, we visualized all summary statistics using violin plots and the GC percentages with box and whisker plots.

We calculated amino acid frequencies and relative synonymous codon usage (RSCU; (Sharp et al. 1986) for each taxon using GCUA (McInerney 1998). Correspondence analyses of amino acid frequencies and RSCU were also performed using GCUA. For each of the first four axes of the correspondence results, we tested for differences between marine and freshwater taxa using the Mann-Whitney U test.

*Ortholog Datasets*

We built three datasets to estimate species trees and explore phylogenomic discordance. The first dataset consisted of the amino acid orthologs, hereafter referred to as the AA dataset. After visualization of GC content variability across first, second, and third codon positions, we excluded all third codon positions from further analysis. We then trimmed and inferred gene trees from the first and second codon position alignments as described above. The second dataset is hereafter referred to as the CDS12 dataset. Next, rapidly evolving synonymous sites are mostly uninformative at deep levels and can lead to long branch attraction (LBA) and compositional heterogeneity (Zwick et al. 2012). To further examine the extent of saturation in the CDS ortholog datasets, we used the degen v.1.4 Perl script (Regier et al. 2010; Zwick et al. 2012) to exclude synonymous signals and recode the full CDS alignments (all three codon positions) using nucleotide ambiguity codes. For example, all leucine codons (CTN, TTR) are degenerated and replaced by YTN. We trimmed and inferred gene trees for each recoded alignment as described above. This third dataset is hereafter referred to as the DEGEN dataset.

We refer to each AA, CDS12, and DEGEN dataset in its entirety as the “complete” datasets. We created two additional subsets of each dataset that sought to maximize phylogenetic information or minimize missing data. First, to maximize signal and reduce stochastic errors, we sorted the complete datasets by the proportion of parsimony-informative sites and subset these to include the top 25% ranked orthologs ("top-PI"). Second, to minimize the amount of missing data and maximize taxon occupancy, we sorted the complete datasets by the number of taxa and subset these to include the top 25% ranked orthologs with highest taxon occupancy ("top-Taxa").

Prior to species tree estimation, we assessed whether dataset partitions violated phylogenetic model assumptions of stationarity, reversibility, and homogeneity (SRH) (Felsenstein 2004). Violations of these assumptions can bias estimates of tree topology and model parameters (Naser-Khdour et al. 2019). We tested each partition in the complete, top-PI, and top-Taxa datasets with the matched-pairs tests of symmetry implemented in IQ-TREE (Naser-Khdour et al. 2019) and excluded partitions with $P < 0.05$.

*Phylogenomic Analyses*

We estimated species trees for each of the complete, top-PI, and top-Taxa datasets using two different approaches: the maximum-likelihood method in IQ-TREE v.2.0.3 using a concatenated data matrix, and the summary quartet method in ASTRAL-III v.5.7.3 (Zhang et al. 2018) using the final estimated ortholog gene trees.

For IQ-TREE, we partitioned each data matrix by gene (ortholog), used ModelFinder to determine the best-fit substitution model for each partition ('-m TEST'), estimated branch support using 10,000 ultrafast bootstrap ('-bb 10000') and 10,000 Shimodaira-Hasegawa-like approximate likelihood ratio test ( '-alrt 10000') replicates (Guindon et al. 2010; Minh et al. 2013), and performed five independent tree searches ('--runs 5'). For ASTRAL, we used the selected ortholog gene trees as input and estimated quadripartition support using local posterior probability (Sayyari and Mirarab 2016).

Standard maximum-likelihood analyses assume a single rate matrix for each partition in a dataset; however, compositional heterogeneity is known to be widespread in phylogenomic datasets. IQ-TREE provides many site-specific frequency models such as the posterior mean site frequency (PMSF; Wang et al. 2018) model as a rapid approximation of the time- and memory-intensive profile mixture models (C10–C60; Si Quang et al. 2008). The PMSF is the amino acid profile for each alignment site computed from an input mixture model and a guide tree. Simulations and empirical studies have demonstrated that PMSF models can be effective against long-branch attraction artifacts (Wang et al. 2018). We estimated a guide tree and species tree in IQ-TREE using the concatenated alignment of the overlapping partitions between the AA-top-PI and AA-top-Taxa datasets ("AA-PMSF"). For the guide tree, we used the LG+F+G model. For the final species, we implemented the PMSF approximation of the LG+C20+F+G model and assessed branch support with 10000 ultrafast bootstrap replicates ('-bb 10000'). The gene trees corresponding to the AA-PMSF dataset were also used as input to ASTRAL.

We compared all 18 species trees using pairwise Robinson-Foulds (RF) distances calculated using the R package *treespace* v.1.1.4.1 (Jombart et al. 2017). We visualized the RF distances using a multidimensional scaling plot. We calculated a majority-rule consensus tree separately for all trees, each method (IQ-TREE, ASTRAL), and each character type (AA, CDS12, DEGEN) using the 'consensus' function in the R package *APE* v.5.6-1 (Paradis et al. 2004).

*Gene Tree and Species Tree Support*

We applied the following approaches to assess the degree of discordance along the species trees. These methods were applied to the species trees estimated from the complete, top-PI, top-Taxa, and PMSF datasets. For these calculations we used the alignments and gene trees of the AA-complete dataset. First, because standard measures of branch support, such as the bootstrap, do not provide a comprehensive view of the underlying agreement or disagreement among genes or sites for a topology, we estimated gene concordance factors (gCF) and site concordance factors (sCF) for each species tree using IQ-TREE (Minh et al. 2020a). For each branch in a species tree, gCF is the number of decisive gene trees that contain that branch, while accounting for variable taxon coverage, and can range from 1 to 100. Meanwhile, sCF is the number of decisive sites that support that branch and generally ranges from ≈33.3 to 100, with 33.3 being the null expectation of no signal. gCF was calculated using the estimated amino acid ortholog trees. sCF was calculated using the concatenated alignments and randomly sampling 1000 sites for each branch in species trees.

Second, as a complement to gCF and sCF, we calculated quartet concordance (QC) factors using QuartetSampling v.1.3.1.b (Pease et al. 2018). For each branch in a species tree, quartets of taxa and gene partitions are randomly sampled and the likelihood of all three possible quartets is evaluated from the sequence data. QC values can range from 1 (all sampled quartets are concordant with the species tree) to 0 (equivalent concordant and discordant quartets) or <0 (more discordant than concordant quartets). For each branch in the species trees, we used the gene-tree mode ('--genetrees *partition-file*') of QuartetSampling to randomly sample 500 ('--reps 500') quartets and partitions, calculated likelihoods using RAxML v.8.2.11 (Stamatakis 2014) ('--engine raxml'), and used a minimum log-likelihood threshold of 2 to evaluate quartet topologies ('--lnlike 2').

*Approximately Unbiased Tests*

To assess the statistical support for each of the five different topologies recovered for the *Thalassiosira* grade (Table 1; Figs. 1–2), we performed Approximately Unbiased (AU) tests (Shimodaira 2002). We tested the support of the AA, CDS12, and DEGEN datasets for each of the five topologies. Constraint trees that represented each of the five topologies were constructed using Mesquite (http://www.mesquiteproject.org). AU tests were implemented in IQ-TREE using the concatenated alignments of the complete datasets, with 10000 RELL replicates (Kishino et al. 1990), and estimating model parameters using an initial parsimony tree.

*Gene Genealogy Interrogation*

To interrogate discordance and the five competing topological hypotheses among the *Thalassiosira* grade, we applied the gene genealogy interrogation procedure (GGI; (Arcila et al. 2017). For this we applied the GGI procedure to the key backbone nodes identified as discordant across the different species tree analyses (nodes A–E; Fig. 1). We tested the five alternative rooted topologies estimated from species tree analyses that only differed in respect to the relationships among the five main lineages of the *Thalassiosira* grade: (1) cyclostephanoids, (2) *Thalassiosira* I, (3) *Thalassiosira* II, (4) *Thalassiosira* III, and (5) *Thalassiosira* IV (Figs. 1–2). We constructed constraint trees using Mesquite and estimated constrained gene trees in IQ-TREE for the top-PI datasets for these five hypotheses ($n_{AA}$=6570, $n_{CDS12}$=3305, $n_{DEGEN}$=3855). The top-PI dataset was chosen because it contains the most signal-rich orthologs. To assess the topology support of each ortholog, we compared site-wise likelihood scores across all five constrained trees with the AU test in IQ-TREE using 10000 RELL replicates. For each ortholog, we ranked the constrained gene trees by AU test *P* value. We then conducted ASTRAL analysis with either: (1) all rank 1 constrained trees, or (2) the set of rank 1 trees significantly better than the alternatives ($P < 0.05$; $n_{AA}$=142, $n_{CDS12}$=56, $n_{DEGEN}$=71; (Arcila et al. 2017).

Despite there being support for the monophyly of the *Thalassiosira* grade clades that we constrained, it is possible that some gene trees show instances of deep coalescence and do not recover the monophyly of some clades. The GGI procedure assumes that all lineages are monophyletic, therefore unsorted allelic polymorphisms can bias the results of the analyses (Arcila et al. 2017). We tested whether the assumption of clade monophyly is met for the *Thalassiosira* grade clades based on coalescent theory predicting that under a neutral coalescent model >99% of genes in a genome will achieve monophyly after 5.3-8.3 coalescence time units (Rosenberg 2003). We used our fossil-calibrated phylogeny (see below) and converted branch lengths in millions of year ($T$) into coalescent units ($\tau$) using the formula $\tau = T / (2N_e$ * generation time). We used a range of estimated effective population sizes that included the estimate for *Phaeodactylum tricornutum* (*Ne*=8.72e6) and values that were twice as large (*Ne*=1.7e7) or twice as small (*Ne*=4.4e6) (Krasovec et al. 2019). We used generation times of 1, 2, or 7 days.

Estimated coalescent unit ($\tau$) branch lengths for *Thalassiosira* grade clades.

| | *Ne* | 4.4e6 | | | 8.72e6 | | | 1.7e7 | | |
|---|---|---|---|---|---|---|---|---|---|---|
| | Gen. time | 1 day | 2 days | 7 days | 1 day | 2 days | 7 days | 1 day | 2 days | 7 days |
| *Thalassiosira* I | | 585.43 | 292.72 | 41.77 | 295.40 | 147.70 | 21.08 | 151.52 | 75.76 | 10.82 |
| *Thalassiosira* II | | 337.72 | 168.86 | 24.10 | 170.41 | 85.20 | 12.16 | 87.41 | 43.70 | 6.24 |
| cyclostephanoids | | 998.12 | 499.06 | 71.22 | 503.64 | 251.82 | 35.94 | 258.34 | 129.17 | 18.43 |
| *Thalassiosira* III | | 267.39 | 133.69 | 19.08 | 134.92 | 67.46 | 9.63 | 69.21 | 34.60 | 4.94 |
| *Thalassiosira* IV | | 455.69 | 227.85 | 32.52 | 229.94 | 114.97 | 16.41 | 117.94 | 58.97 | 8.42 |

*Polytomy Tests*

To test if any branches in the species trees might be better represented as a polytomy, we performed the polytomy test implemented in ASTRAL (Sayyari and Mirarab 2018). We used the gene trees from the complete datasets, collapsing low supported branches with ultrafast bootstrap support <33 or <75. Results for each branch were significant at $P < 0.05$ to reject the null hypothesis of a polytomy.

*Divergence Time Analysis*

The PMSF species tree was chosen as the reference tree for estimation of divergence times using the Bayesian approach in MCMCtree v.4.9e (Rannala and Yang 2007; Yang 2007). For this analysis, we pruned the tree of outgroup taxa. We selected orthologs from the AA-PMSF dataset to use for the dating analysis using two criteria: (1) the degree of violation from the molecular clock (DVMC) and (2) the normalized Robinson-Foulds (nRF) distance from the reference PMSF tree. Lower values of DVMC are considered desirable as they are indicative of a lower degree of violation in the molecular clock assumption. Lower values of nRF distance are considered acceptable as they reflect a greater similarity of the ortholog tree to the species tree. We calculated DVMC and nRF using PhyKit and selected orthologs with DVMC <0.3 and nRF <0.3 to include in the dating analysis, resulting in 79 orthologs. We then used PartitionFinder (Lanfear et al. 2016) as implemented in IQ-TREE to merge the alignments of the 79 ortholog partitions and estimate the best-fit substitution model for each merged partition. This approach merged the 79 partitions into 19 partitions and identified the Jones-Taylor-Thornton (JTT) model as the best-fit model for each partition.

We applied the following five constraints and fossil calibrations to the tree based on previous divergence time estimates conducted in (Alverson 2014; Nakov et al. 2018).

1. Crown Thalassiosirales–Lower bound of 75 Ma based on the *Thalassiosiropsis* fossil used in (Alverson 2014). Upper bound of 118 Ma based on the estimate in (Nakov et al. 2018). Constraint=”B(0.75,1.18)”.
2. *Porosira glacialis*–Lower bound of 9 Ma based on the first appearance in the Neptune database (Lazarus 1994). Constraint=”L(0.09,0.1,1)”.
3. *Cyclostephanos*–Lower bound of 5 Ma based on the fossil *Cs. undatus (Theriot and Kociolek 1986; Krebs and Kociolek 1994)*. Constraint=”L(0.05,0.1,1)”.
4. *Bacterosira constricta*–Lower bound of 8.35 Ma based on the first appearance in the Neptune database (Lazarus 1994). Constraint=”L(0.0835,0.1,1)”.
5. *Shionodiscus*–Lower bound of 6.15 Ma based on the fossil *Sh. praeoestrupii (Shiono and Koizumi 2000)*. Constraint=”L(0.0615,0.1,2)”.

We applied the approximate likelihood calculation for divergence time estimation in MCMCtree (Reis and Yang 2011). For each of the 19 partitions, we estimated the branch lengths, gradient, and Hessian using CODEML with the JTT+G5 model. We then combined each Hessian and supplied it to MCMCtree (usedata=2). We assigned the Dirichlet-gamma prior for the overall substitution rate as G(2,20) with a mean of 1 corresponding to $10^{-8}$ substitutions per site per year (rgene_gamma=2 20 1), the Dirichlet-gamma prior for the rate-drift parameter as G(1,10) (sigma2_gamma=1 10 1), and the parameters for the birth-death sampling process with birth and death rates $\lambda=\mu=1$ and sampling fraction $\rho=0.1$ (BDparas=1 1 0.1). We estimated divergence times using the autocorrelated relaxed rates model (clock=3) and ran the MCMC chain for 10 million generations (nsample=3000, sampfreq=250), discarding the first 2.5 million generations as burnin (burnin=2500000). Two independent runs of each rate model were performed to ensure model convergence. MCMC chains were summarized and estimated sample sizes (ESS) for all parameters were calculated using Tracer v.1.7.2 (Rambaut et al. 2018). We plotted the estimated divergence times using the R package *MCMCtreeR* v.1.1 (Puttick 2019).

*Ancestral State Reconstruction of Habitat*

We used the dated species tree for analyses of trait evolution, but we downsampled the tree by removing all conspecific strains and leaving one tip per species. For each species, we scored whether the species was found in marine (state 0) or freshwater (state 1) habitats. Euryhaline taxa were assigned to the marine state (state 0). Using the dated phylogeny and character states, we performed marginal ancestral state reconstruction using the SSE framework (state-dependent speciation and extinction; Maddison et al. 2007; FitzJohn et al. 2009; Beaulieu and O'Meara 2016). Our tree included 53 of 580 extant species as estimated from AlgaeBase (Guiry and Guiry 2022). To correct for incomplete taxon sampling, we estimated the proportion of sampled taxa in each habitat by dividing the number of species in the tree by the total number of extant species found in each habitat. For the SSE approach, we compared five models: (1) a "trivial null" model with one parameter for the turnover rate for both observed states and one parameter controlling the rate of character transitions; (2) a BiSSE model (binary-state SSE; (FitzJohn et al. 2009) with separate turnover rates estimated for lineages living in the different habitats and asymmetric transitions between habitats; (3) a HiSSE model (hidden-state SSE; (Beaulieu and O'Meara 2016) with separate turnover rates for each combination of observed and hidden states and eight transition rates for changes between habitats; (4) a CID–2 model (character-independent; (Caetano et al. 2018) with two turnover rate parameters similar to the BiSSE model, but not linked to the observed character states; and (5) a CID–4 model with five diversification parameters matching the complexity of the HiSSE model (Caetano et al. 2018). Each model had a single extinction fraction parameter (defined as extinction/speciation, which measures the ratio of death and birth per unit time) kept constant across states. We compared the fit of all five models using the Akaike information criterion (AIC; Akaike 1998). We performed marginal ancestral state reconstruction using the parameters estimated in the best-fit model. All SSE analyses were conducted in the R package *hisse* v.2.1.6 (Beaulieu and O'Meara 2016).

Standard methods of ancestral state reconstruction use only a single estimate of species relationships to estimate model parameters. However, extensive gene tree-species tree discordance can lead to incorrect inferences of trait evolution if only a single species tree is used for analyses. This phenomenon has been termed hemiplasy (Avise and Robinson 2008; Hahn and Nakhleh 2016) and is an important consideration in rapidly radiating groups. We explored the potential for hemiplasy in explaining the origin of freshwater taxa in the Thalassiosirales. We estimated the Hemiplasy Risk Factor (HRF) for each internal branch of the species tree using the R package *pepo* v.0.2 (Guerrero and Hahn 2018). The HRF represents a way to describe the relative probabilities of hemiplasy and homoplasy for an incongruent trait. We explored how differences in effective population size ($N_e$) and generation time affect HRFs. We converted branch lengths in millions of year ($T$) into coalescent units ($\tau$) using the formula $\tau = T / (2N_e *$ generation time$)$. We estimated nine sets of coalescent branch lengths using combinations of three $N_e$ estimates (4.35e6, 8.72e6, 1.74e7) and four generation times (1 day, 7 days, 1 month, 6 months). For calculating HRF per branch, we supplied *pepo* with the branches lengths in coalescent units and a population-wide mutation rate estimate, $\Theta$=0.0125. The mutation rate estimate is taken from the diatom *Phaeodactylum tricornutum,* which to date has been the only diatom with an experimentally estimated mutation rate (Krasovec et al. 2019).

*Summarizing marine–freshwater character histories*

We used a simple parsimony-based approach to count and summarize the number of habitat character histories that potentially involve hemiplasy. We chose the set of orthologs that contained outgroup taxa (n=4811). For each ortholog we used both the estimated gene tree topology (GT) as well as a topology constrained to the PMSF species tree (ST). We estimated branch lengths for each constrained ST using IQ-TREE with the JTT+F+I+G model. For each ortholog, we rooted both GT and ST on an available outgroup taxon, since not all outgroups are present in each ortholog. We used the *pxrr* command from Phyx (Brown et al. 2017) and supplied a ranked list of outgroup taxa, which was used to root each tree at the first available outgroup from the ranked list. We coded all taxa in the GT and ST for habitat (0=marine; 1=freshwater). Since all outgroup taxa are marine, we do not expect inexact outgroup rooting to affect reconstructions. We performed separate parsimony optimization of habitat using both the GT and ST topologies for each ortholog under the accelerated transformation (ACCTRAN) method employed in the R package *paleotree* (Bapst 2012). ACCTRAN assigns changes closer to the root and attempts to minimize the number of parallel origins for complex traits, whereas alternative methods (such as maximum parsimony reconstruction [MPR]) assign changes as close to the tips as possible (Agnarsson and Miller 2008). We ran both ACCTRAN and MPR methods and saw few differences. We then tabulated the minimum number of reconstructed step transitions from marine to freshwater (0→1) on both GT and ST topologies. Similar to (Copetti et al. 2017), we calculated the difference between the number of transitions optimized on the GT and ST topologies. We placed each ortholog into one of three groups: (1) the ortholog had fewer transitions on the GT than the ST (GT < ST); (2) the ortholog had no difference in the number of transitions between the ST and GT (GT = ST); or (3) the ortholog had more transitions on the GT than the ST (GT > ST). Orthologs in the first group (GT < ST) are those that have at least one hemiplasy.

*Ancestral sequence reconstruction*

We quantified the number of amino acid sites displaying signatures of hemiplasy or homoplasy using ancestral sequence reconstruction. We used the same set of orthologs as above (see *Summarizing character histories*). Prior to the analyses, we trimmed the amino acid alignments to remove large gaps (≥95% column occupancy), while retaining parsimony-informative sites and constant sites. For each ortholog, we used the trimmed alignment to reconstruct ancestral sequences at each tree node using both the GT and ST topologies using PAML (Yang 2007). By default, PAML uses unrooted trees during analyses. For reconstruction, we used fixed branch lengths (previously inferred for both GT and ST) and used the JTT+F+G model since PAML does not implement invariant site models. For each alignment site in each ortholog, we tabulated the number of amino acid transitions that were estimated on both the GT and ST topologies and calculated the difference (GT minus ST). Similar to above, the sites were placed into one of three groups: (1) the site had fewer transitions on the GT than the ST (GT < ST); (2) the site had no difference in the number of transitions between the ST and GT (GT = ST); or (3) the site had more transitions on the GT than the ST (GT > ST). Orthologs in the first group (GT < ST) are those that have at least hemiplasy.

*Ortholog comparison to* Skeletonema marinoi

To determine whether the set of orthologs containing hemiplasies (n = 647) might be related to salinity adaptation, we compared them with differentially expressed (DE) genes from a marine/brackish salinity experiment in *Skeletonema marinoi (Pinseel et al. 2022)*. We used NCBI BLASTp to identify matching protein sequences between *Sk. marinoi* and the set of orthologs containing hemiplasies. Functional annotation of the *Sk. marinoi* gene models and proteins are described in (Pinseel et al. 2022).

*Tests for Recombination*

We tested for signatures of intralocus recombination using the full coding sequence alignments (all three positions) with PhiPack (Bruen et al. 2006). We used PhiPack to calculate the pairwise homoplasy index (PHI) with the default sliding window of 100 bp. An alignment was considered recombinant if it had a Bonferroni-adjusted $P$ value < 0.05.

*Data Availability*

Newly generated sequencing and assembly data have been deposited in NCBI BioProject PRJNA825288. Sequencing reads have been deposited in the Sequence Read Archive (SRA). Whole Genome Shotgun (WGS) and Transcriptome Shotgun Assembly (TSA) projects have been deposited at GenBank. SRA, WGS, and TSA accession numbers are listed in Supplementary Table 2. Alignments, gene trees, species trees, and scripts are available from: https://doi.org/10.5061/dryad.7m0cfxpxp. Voucher images, proteomes, alignments, trees, log files, and code have been deposited in Zenodo: https://doi.org/10.5281/zenodo.7713227.

UniProt Consortium. 2021. UniProt: the universal protein knowledgebase in 2021. Nucleic Acids

Res. 49:D480–D489.

Wang H.-C., Minh B.Q., Susko E., Roger A.J. 2018. Modeling Site Heterogeneity with Posterior Mean Site Frequency Profiles Accelerates Accurate Phylogenomic Estimation. Syst. Biol. 67:216–235.

Yang Y., Smith S.A. 2013. Optimizing de novo assembly of short-read RNA-seq data for phylogenomics. BMC Genomics. 14:328.

Yang Y., Smith S.A. 2014. Orthology inference in nonmodel organisms using transcriptomes and low-coverage genomes: improving accuracy and matrix occupancy for phylogenomics. Mol. Biol. Evol. 31:3081–3092.

Yang Z. 2007. PAML 4: phylogenetic analysis by maximum likelihood. Mol. Biol. Evol. 24:1586–1591.

Zhang C., Rabiee M., Sayyari E., Mirarab S. 2018. ASTRAL-III: polynomial time species tree reconstruction from partially resolved gene trees. BMC Bioinformatics. 19:153.

Zwick A., Regier J.C., Zwickl D.J. 2012. Resolving discrepancy between nucleotides and amino acids in deep-level arthropod phylogenomics: differentiating serine codons in 21-amino-acid models. PLoS One. 7:e47450.
