## Supplementary Materials description for "Resolving marine–freshwater transitions by diatoms through a fog of discordant gene trees"

Note on taxon labels:

- Genomes have strain ID before genus/species (e.g., CCMP332_Cyclotella_cryptica).
- Transcriptomes have strain ID after genus/species (e.g., Cyclotella_nana_AJA048-54).

**Supplementary Figure S1**. Violin plots showing the Relative Composition Variability (RCV) for amino acids (AA), first codon positions (CDS-pos1), second codon positions (CDS-pos2), and third codon positions (CDS-pos3). RCV values range from 0 to 1, with lower values signifying less compositional heterogeneity.

**Supplementary Figure S2**. GC content variability across taxa and orthologs for first (left), second (center), and third codon positions (right). Marine taxa are colored yellow, freshwater taxa are colored purple. Taxa are ordered within each subplot by median GC content.

**Supplementary Figure S3.** Violin plots showing the estimated saturation for amino acids (AA), complete coding sequences (CDS), first and second codon positions (CDS12), and degenerate codon sequences (DEGEN). Saturation is calculated for each ortholog (n=6262) as the regression slope between the branch length patristic distances and uncorrected sequence distances.

**Supplementary Figure S4**. Biplots showing the results of correspondence analyses for relative synonymous codon usage (top left) and amino acid frequencies (bottom left). The first two dimensions from correspondence analysis are shown as Axis 1 and Axis 2. Marine taxa are colored green, freshwater taxa are colored blue. Unique shapes correspond to taxa in several named clades (see Figure 1). Mann-Whitney U test results for Axis 2 (top right; bottom right) show separation between marine and freshwater taxa.

**Supplementary Figure S5**. Multidimensional scaling plot of pairwise Robinson-Foulds distances between all species trees. Labels for each point note the character type and data subset that were used for that species tree. Points are colored based on the species tree methodology. Shapes correspond to the returned topology for the *Thalassiosira* grade.

**Supplementary Figure S6**. Gene (gCF; left) and site (sCF; right) concordance factors estimated using the AA-complete dataset and the PMSF species tree. CF=concordance factor (purple); DF1=discordance factor of first nearest-neighbor exchange (green); DF2=discordance factor of second nearest-neighbor exchange (yellow); DFP=discordance due to polyphyly (grey).

**Supplementary Figure S7**. Quartet concordance factors (QC) estimated on the PMSF species tree using the AA-complete (left) and CDS12-complete (right) datasets. Branch colors denote relative QC support: moderate to strong support for the focal branch (QC > 0.2); weak support for the focal branch ($0 < QC <= 0.2$); weak support for an alternative quartet ($-0.05 < QC <= 0$); moderate to strong support for an alternative quartet ($QC <= -0.05$).

**Supplementary Figure S8**. Comparison of concordance factors between the AA-complete and AA-top-PI datasets on the PMSF species tree. Gene concordance factors (gCF) are shown above each branch. Site concordance factors (sCF) are shown below each branch.

**Supplementary Figure S9**. Comparison of concordance factors and estimated node ages. Node ages are in millions of years (Ma) estimated by MCMCtree. Gene concordance factors (gCF) are shown in blue. Site concordance factors (sCF) are shown in orange. The blue and orange lines indicate the regression line of CF ~ node age and include the slope intersect formula and $R^2$ values.

**Supplementary Figure S10**. Divergence time estimates for the Thalassiosirales using MCMCtree. Node ages are summarized in a million year timescale. Node age uncertainty is shown with blue bars and represents the 95% credibility interval of estimated times. Geologic period and epoch boundaries are shown below the tree and highlighted with gray backgrounds behind the tree. Ea.=Early Cretaceous; La.=Late Cretaceous; P.=Paleocene; Eo.=Eocene; Mi.=Miocene; Pa.=Paleogene; Ne.=Neogene.

**Supplementary Figure S11.** Hemiplasy Risk Factors calculated using a range of effective population sizes (Ne) and generation times. Branches are colored based on their calculated HRF. Scale bars represent the branch length in estimated coalescent time units.

**Supplementary File S1**. Materials and methods.

**Supplementary Table S1**. Strain collection and voucher information.

**Supplementary Table S2**. Assembly accession numbers and proteome information.

**Supplementary Table S3**. Ortholog summary statistics.

**Supplementary Table S4**. Approximately unbiased test results for the *Thalassiosira* grade.

**Supplementary Table S5**. Gene genealogy interrogation results for the *Thalassiosira* grade.

**Supplementary Table S6**. Results of HiSSE model testing.

**Supplementary Table S7**. Summary of hemiplasy scenarios within the highest taxon occupancy subset of hemiplasy orthologs.

**Supplementary Table S8.** Functional annotations of the hemiplasy orthologs with BLAST hits to proteins in the *Skeletonema marinoi* genome.
